# Novel carboxy-terminal Pannexin 1 isoform is expressed in mouse melanoma cells and interacts with the full-length protein

**DOI:** 10.1101/2024.09.09.612143

**Authors:** Brooke L. O’Donnell, Dan Stefan, Yu-Hsin Chiu, Michael J. Zeitz, Stephanie E. Leighton, Laura A. Orofiamma, Danielle Johnston, Justin Tang, Carlijn Van Kessel, Kevin Barr, Laszlo Gyenis, Taylor J. Freeman, John J. Kelly, Samar Sayedyahossein, Brant E. Isakson, David W. Litchfield, Kathryn Roth, James W. Smyth, Matthew Hebb, John Ronald, Douglas A. Bayliss, Silvia Penuela

## Abstract

Alternative translation initiation (ATI) is a process of increasing protein diversity from one transcript, allowing cells to rapidly respond to signals, which is particularly important in cancer cells. Here, we report potential internal translation start sites exist in PANX1 which have implications in trafficking and channel function. Using mouse (mPANX1) constructs for each internal methionine, we saw that these PANX1 isoforms were N-glycosylated, could traffic to the cell surface and mPANX1-M37 formed functional channels activated by C-terminus cleavage or α1-adrenoceptor stimulation. We also identified a ∼25 kDa isoform of mPANX1 (mPANX1-25K) endogenously expressed in mouse melanoma cell lines that could be confirmed with a cognate peptide. mPANX1-25K lacks the mPANX1 N-terminus and most likely corresponds to the M210 internal translation start site since we could not identify any alternative transcripts that would produce this ATI product. When we expressed the human equivalent of M210 in Hs578T *PANX1* KO cells with and without wildtype human PANX1, we determined M211 exhibits a predominantly intracellular localization, is N-glycosylated and can interact with full-length human PANX1. Collectively, these findings indicate species specific differences in the abundance of PANX1 ATI isoforms which could act independently or in conjunction with the canonical full-length protein in melanoma.

**Significance Statement:**

- Alternative translation initiation increases the number of proteins produced from a single mRNA. Connexin internal translation isoforms have been characterized in a wide variety of cell contexts and diseases, but it is unknown whether the related pannexins also undergo this process.
- We discovered a novel 25 kDa mouse pannexin 1 protein (mPANX1-25K), most likely corresponding to an internal methionine start site at M210, which is N-glycosylated and intracellularly localized. This isoform is not expressed in humans.
- This represents the first report of a PANX1 alternative translation initiation isoform and points to an evolutionary divergence of mouse and human PANX1.

## INTRODUCTION

Alternative translation initiation (ATI) is any non-canonical translation that occurs independently of the 5’ 7-methylguanosine cap, eukaryotic initiation factor 4F binding complex and the first AUG (methionine) codon present in an mRNA transcript. ATI, which was initially thought to only occur in viruses, has been increasingly studied in eukaryotic cells where it acts to expand protein diversity by increasing the number of proteins generated from one transcript. ATI also provides an avenue for cells to rapidly respond to cellular signals and stresses using transcripts already present within the cell. This process has been shown to function as either an adaptation to stress or contributor to diseases like cancer (Kozak, 1992; Kochetov, 2008; James and Smyth, 2018; Lacerda *et al*., 2019). Many modes of ATI have been reported which are often favoured in times when cap-dependent translation is compromised or inhibited, including internal ribosome entry sites, ribosome shunting and leaky ribosome scanning (Komar and Hatzoglou, 2011; James and Smyth, 2018; Kwan and Thompson, 2019).

Connexin 43 (Cx43, *GJA1*), a member of the gap junction protein family, has been shown to produce several ATI variants corresponding to six internal, in-frame methionine codons in several different cell and tissue types. Cx43 C-terminal (CT) isoforms include GJA1-32k (M100), GJA1-29k (M125), GJA1-26k (M147), GJA1-20k (M213), GJA1-11k (M281) and GJA1-7k (M320) which were initially reported in human heart lysates (Smyth and Shaw, 2013; Epifantseva and Shaw, 2018). However, analysis of the function of GJA1-20k has largely been the focus of subsequent research (Fournier *et al*., 2025), where this shorter isoform autoregulates Cx43 hexamer oligomerization, trafficking and gap junction formation (Smyth and Shaw, 2013; Salat-Canela *et al*., 2014). Without GJA1-20k, the half-life of cytoplasmic Cx43 is halved due to rapid degradation triggered by improper trafficking (Xiao *et al*., 2020). In disease states such as chronic ischemia and ischemia reperfusion injury, GJA1-20k has cardioprotective effects when overexpressed (Basheer *et al*., 2017; Basheer *et al*., 2018) but is downregulated in aged hearts as well as hypoxic and transforming growth factor-β-treated cells (Kotini *et al*., 2018; Zeitz *et al*., 2019). *GJA1^M213L/M213L^* mice, which are homozygous for a mutation to prevent the translation of GJA1-20k, are not viable past 2-4 weeks of age, with sudden death due to abnormal cardiac electrical excitation (Xiao *et al*., 2020). Furthermore, GJA1-20k is expressed in prostate, lung, breast, cervical and carcinoma cell lines (Fournier *et al*., 2025), and is downregulated during epithelial to mesenchymal transition, during which cells must break cell-cell adhesions to facilitate migration (James and Smyth, 2018). Overall, this indicates GJA1-20k plays critical roles in normal tissue homeostasis and its levels may be dysregulated in cases of aging, disease or cellular stress, but whether the related pannexin genes are also capable of internal translation has not been investigated.

Pannexins (PANX1, PANX2, PANX3) are heptameric, channel-forming glycoproteins of which PANX1 has been the most extensively studied in tissue development as well as the initiation and progression of various skin, musculoskeletal, cardiovascular, nervous and inflammatory diseases (Yeung *et al*., 2020; Laird and Penuela, 2021; O’Donnell and Penuela, 2021; Rusiecka *et al*., 2022; O’Donnell and Penuela, 2023; Luo *et al*., 2024; O’Donnell *et al*., 2025b). While the canonical function of PANX1 is to act as an ion and metabolite releasing channel at both the cell surface and endoplasmic reticulum, channel-independent functions for PANX1 have also emerged more recently. For example, PANX1 directly binds to the Wnt pathway effector β-catenin in melanoma cells, enabling PANX1-mediated stabilization of β-catenin to regulate melanoma cell growth and mitochondrial respiratory activity (Sayedyahossein *et al*., 2021).

In addition to the full length PANX1 transcript/protein, pannexin isoform transcripts generated through alternative splicing or truncations have been identified in rodents and humans, but few have been detected at the protein level (O’Donnell and Penuela, 2023). RefSeq *PANX1* (O’Leary *et al*., 2016)—also annotated as PANX1b (Ma *et al*., 2009) and PANX1-201 in Ensembl (Cunningham *et al*., 2022)—contains 5 exons and 4 introns that can be alternatively spliced at exon 2, 4 (Li *et al*., 2011) and 5 (Baranova *et al*., 2004a) to produce PANX1c, PANX1d and PANX1a transcript isoforms, respectively. Another PANX1 splice variant which lacks exon 3 was detected in the reproductive tract of male rats, where its abundance may be regulated by androgens (Turmel *et al*., 2011). In highly metastatic breast cancer cells, a shorter version of PANX1 was identified which results from a nonsense mutation such that the protein contains only the first 89 amino acids (PANX1^1-89^). Through augmented ATP release and downstream purinergic signalling, the PANX1^1-89^ mutant promotes the survival of metastatic cells during the process of extravasation at a secondary microvasculature site, where cells would otherwise die from intravascular deformation of the plasma membrane (Furlow *et al*., 2015; Wang *et al*., 2025). Lastly, a translationally incompetent *PANX1* transcript variant which lacks the 5’ untranslated region was identified in rhabdomyosarcoma cells, resulting in a reduction in PANX1 levels in rhabdomyosarcoma cells compared to normal skeletal muscle myoblasts despite containing similar amounts of *PANX1* mRNA (Xiang *et al*., 2022). Because the pannexin family only contains three genes, alternative splicing is one way to increase pannexin protein diversity in different cell types. These pannexin isoforms may have distinct cellular consequences such as altered channel activity and signalling. However, it is currently unknown whether PANX1 can also undergo ATI as well as if these ATI isoforms have specific cellular functions.

In this study, we report the presence of ATI sites in mouse PANX1 (mPANX1) and human PANX1 (hPANX1) and the initial identification of a novel 25 kDa isoform seen in mouse melanoma cells. Using plasmids for the internal methionine start sites, we also characterize PANX1 ATI products glycosylation, trafficking and channel function, as well as their ability to interact with the full-length PANX1 protein.

## RESULTS

### Possible ATI sites are present in both *mPANX1* and *hPANX1*

By analyzing the mouse *mPANX1* transcript sequence, we found 4 additional in-frame AUG methionine codons downstream of the canonical start site at M1 that produces full-length mPANX1 (mPANX1-FL), including M37, M143, M210 and M374 (Figure 1A). Each additional start site is present in the intracellular facing domains of mPANX1, including the N-terminus (NT), intracellular loop and CT. Consistent with the possibility of multiple ATI sites, we observed a range of HA-specific bands (from ∼30-60 kDa) in HEK293T cells expressing mPANX1 with a CT HA tag (mPANX1-HA) via Western blotting (Figure 1B). While complex N-glycosylation at N254 of mPANX1 could lead to multiple immunoreactive bands (Boassa *et al*., 2007; Penuela *et al*., 2007), the deglycosylated samples treated by Peptide-*N*-Glycosidase F (PNGase F) showed a similar, but shifted banding pattern (now ranging from ∼25-50 kDa) instead of collapsing to one non-glycosylated form, indicating that the banding pattern was not due to differential glycosylation. In congruence with the possibility of multiple in-frame isoforms, each internal methionine is surrounded with the Kozak initiation consensus sequence (RxxAUGG; R=A/G) crucial for translation initiation (Kozak, 1986) (Figure 1C). To examine whether the identified AUG codons could function as potential start sites, we generated a series of mutant mPANX1 constructs where we retained only one methionine codon and included isoleucine substitutions at all other sites to markedly reduce translation efficiency (Figure 1D). We found a single band in blots from HEK293T cells expressing HA-tagged mPANX1 constructs that retained only a single ATI site. The apparent molecular weights of these HA antibody-detected signals were consistent with those estimated from the predicted ATI sites, further verifying that M37, M143 and M210 can serve as translation initiation sites alternative to the conventional M1. Notably, no protein expression of the M374-only form of mPANX1 was detectable in HEK293T cells following transient transfections.

**Figure 1.**
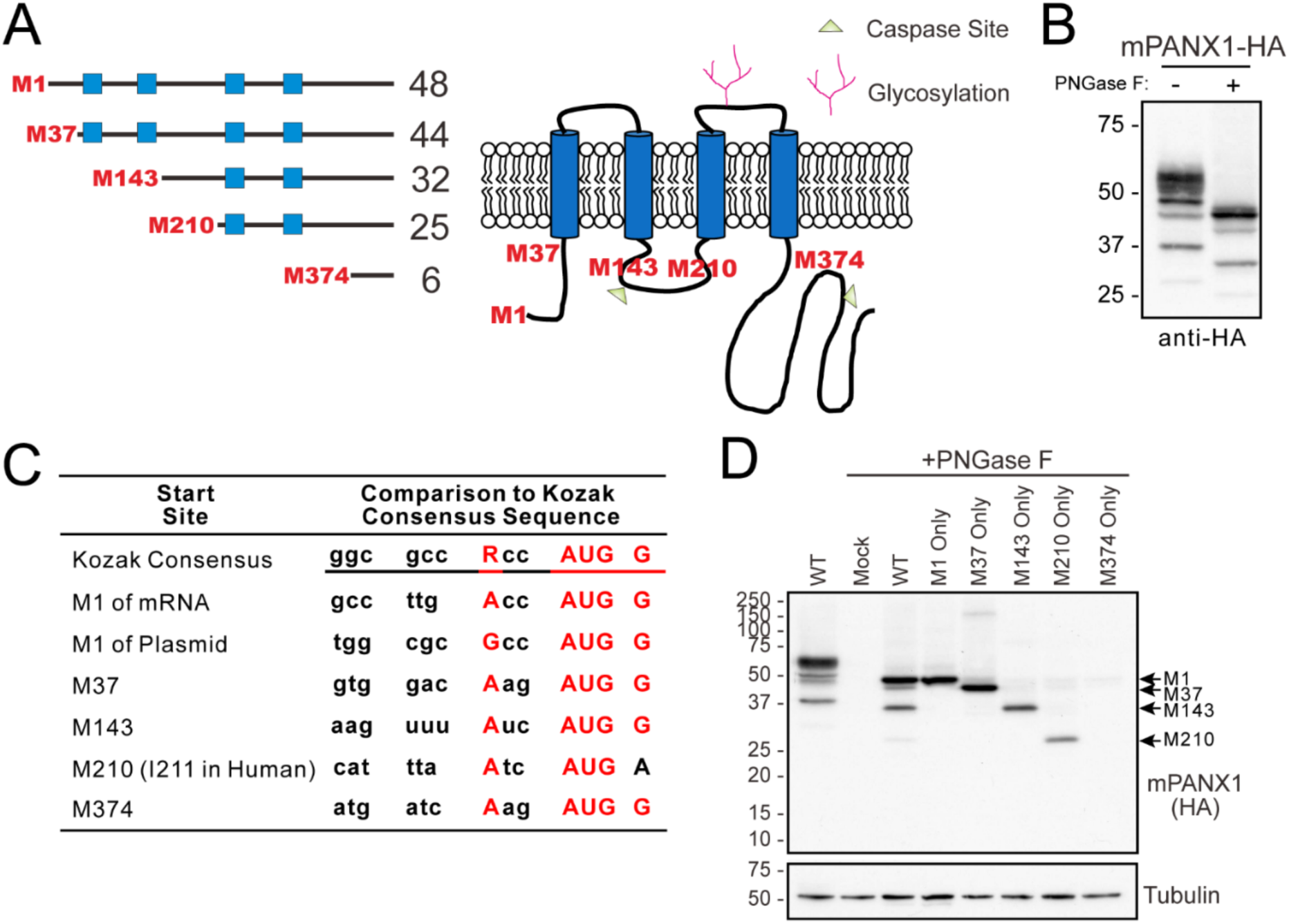
Multiple glycosylated forms of mPANX1 proteins are expressed from different internal translational initiation sites. (A) Schematic depicts predicted molecular weights (in kDa), topology and post-translational modifications of potential mPANX1 ATI isoforms starting at each internal methionine (M) residue. (B) Western blots of HEK293T cells transiently expressing HA-tagged mPANX1 (mPANX1-HA). (C) Sequence alignments showed the *mPANX1* transcript encodes multiple in-frame methionine residues, including M1, M37, M143, M210 and M374, that follow the Kozak initiation consensus sequence (RxxAUGG; R=A/G). Key residues matching the Kozak consensus sequence (first row) are shown in red. (D) Western blotting shows HEK293T cells transfected with wildtype mPANX1-HA (WT), shorter PANX1 isoforms from each internal start site (M1, M37, M143, M210, M374 Only) or empty vector (Mock) constructs and subjected to PNGase F treatment. Arrows indicate varying forms of mPANX1 proteins. M374 was not detected. Tubulin used as protein loading control, protein sizes in kDa.

As with mPANX1, we discovered additional amino acid residues throughout hPANX1 which could function as possible sites of internal translation (Figure 2A, B), including nine internal methionine residues and one isoleucine residue (I211, corresponding to M210 in mice). Like the potential mPANX1 ATI sites, all possible hPANX1 internal start sites are retained in the intracellular domains of hPANX1, with one in the NT, three in the intracellular loop and six in the CT, with predicted molecular weights ranging from 2.8 kDa to 44 kDa. The potential M37, M143, M165 and I211 isoforms retain multiple transmembrane domains—indicating they could be integral membrane proteins—as well as many hPANX1 post-translational modification sites. Analysis of the canonical *hPANX1* transcript for the Kozak initiation sequence at each potential internal start site (Figure 2C) revealed moderate to high sequence similarity to the consensus sequence at most sites. Additionally, exogenous expression of wildtype (WT) hPANX1 in Hs578T *hPANX1* knockout (KO) cells presented four immunoreactive species in addition to the canonical full-length hPANX1 (hPANX1-FL) that were detectable by the PANX1 CT-412 antibody (targeting the end of the CT including residues 412-426 (Penuela *et al*., 2007)) and appeared to have lower molecular weights than hPANX1-FL bands (Figure 2D), resembling some of those observed with mPANX1-HA (Figure 1B). Interestingly, I211 shows evolutionary conservation in most primate species, whereas rodents have either a methionine or valine at the corresponding amino acid position 210 (Figure 3). Both isoleucine and valine have been previously shown to act as non-canonical ATI start sites (Touriol *et al*., 2003; Ingolia *et al*., 2011; Lee *et al*., 2012; Kearse and Wilusz, 2017). Unlike I211, M37, M143 and M375 (M374 in mice) are evolutionarily conserved in all primates and rodent species we compared, indicating potentially strong selective pressure to retain these sites. Apart from M388 which encodes a threonine in a majority of the primate species, the other internal methionine sites present in hPANX1 (M165, M384, M388 and M402) are largely conserved in primates, but differ to a greater degree in rodent species.

**Figure 2.**
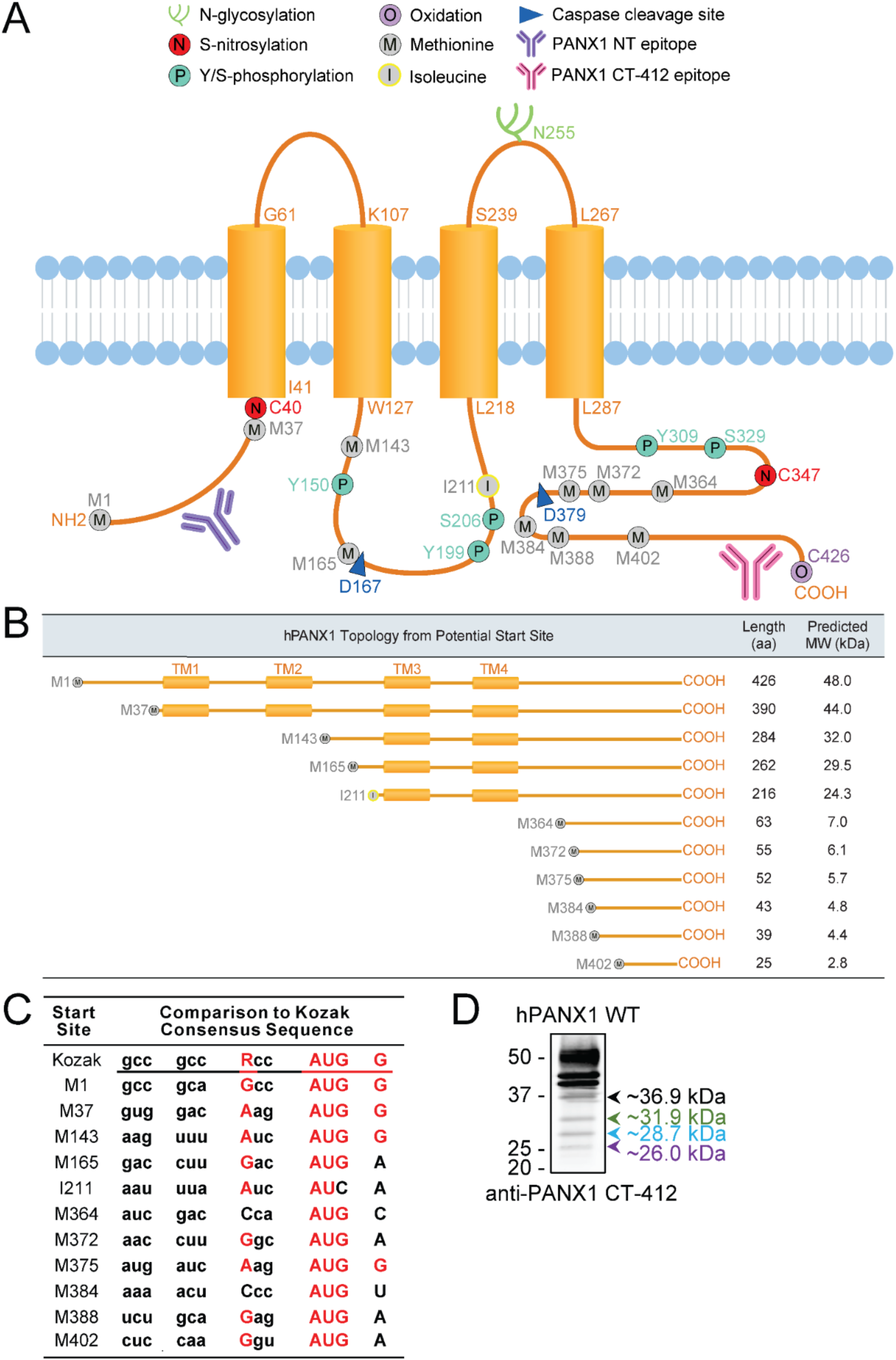
hPANX1 structure and potential internal translation start sites. (A) Schematic depicts hPANX1 topology, post-translational modifications, sites of antibody binding and internal methionine (M) and isoleucine (I) amino acid residues which could act as potential internal start sites. (B) Table depicts hPANX1 topology, amino acid (aa) length and predicted molecular weight in kDa for potential ATI isoforms. Created using Adobe Illustrator CS6. (C) Kozak consensus sequence assessment for potential internal translation start sites of *hPANX1* mRNA shows multiple in-frame methionine and one isoleucine that follow the Kozak initiation consensus sequence (RxxAUGG; R=A/G). Key residues matching the Kozak consensus sequence (first row) are shown in red. (D) Immunoblotting of Hs578T *hPANX1* KO cells expressing wildtype (WT) hPANX1. Blot probed with PANX1 CT-412 antibody.

**Figure 3.**
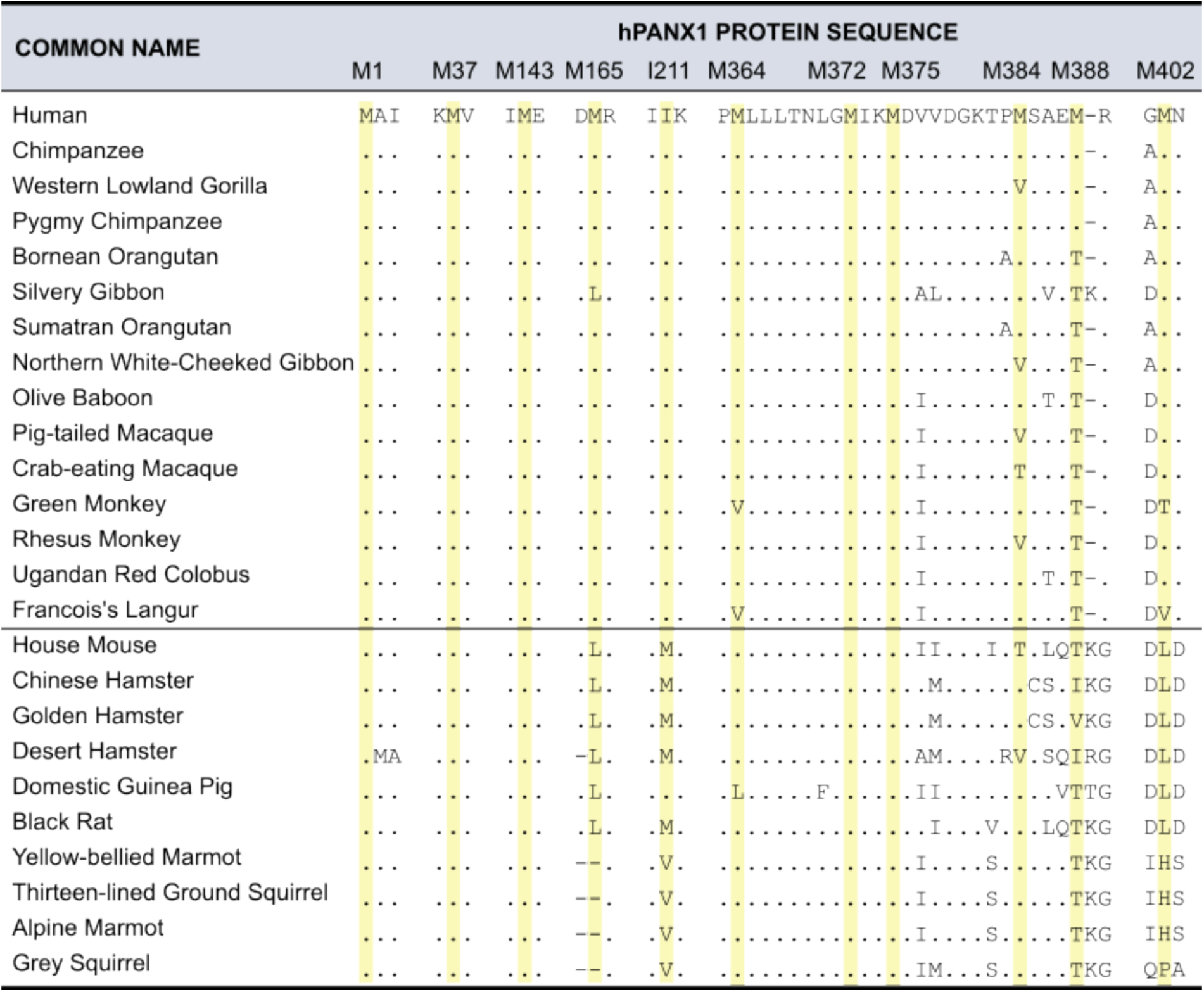
The isoleucine at residue 211 seen in hPANX1 is conserved in most primate species, whereas rodents contain a methionine or valine at the corresponding position. Protein sequence homology was analyzed using NCBI standard protein BLAST. Aligned sequences were compared between hPANX1 and relevant organisms (primates above dividing line, rodents below) with available complete PANX1 sequences. Yellow highlight denotes locations of internal hPANX1 methionines and isoleucine 211 residue and corresponding residues in each organism. Dots represent conserved residues. In the rodent species, dashes represent deletions relative to the human PANX1 sequence and in human and primate PANX1, dashes represent an insertion in the rodent species relative to human/primate.

### A 25 kDa isoform of PANX1 is endogenously present in mouse melanoma cell lines

To determine if any ATI isoforms of PANX1 are present endogenously, we first analyzed isogenic mouse melanoma cell lines B16-F0, -F10 and -BL6 (Figure 4). Immunoblotting with the PANX1 CT-412 antibody revealed a 25 kDa species (mPANX1-25K) which could be significantly competed with peptide pre-adsorption similar to mPANX1-FL. Furthermore, the 25 kDa species was not detectable with a PANX1 NT antibody, suggesting this PANX1 isoform could arise from internal translation since it would lack the NT domain. mPANX1-25K most likely corresponds to the M210 start site based on both the predicted (Figure 1A) and observed (Figure 1D) sizes of the M210 ATI isoform. In this case, mPANX1-25K would contain part of the intracellular loop, second extracellular loop and the complete CT of mPANX1-FL.

**Figure 4.**
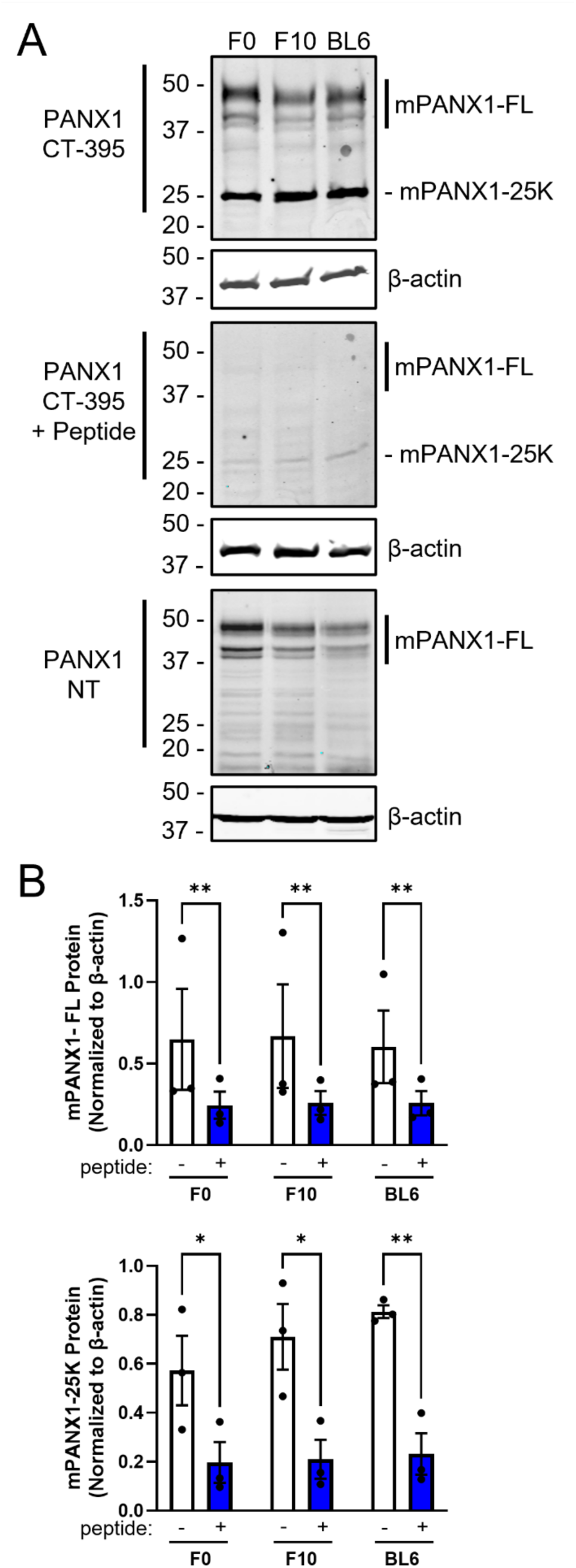
Endogenous mPANX1-25K is present in mouse cell lines. (A) Western blot of B16-F0, F10 and BL6 melanoma cell lines with the mPANX1 C-terminal (CT)-395 antibody with and without peptide competition as well as the PANX1 N-terminal (NT) antibody. β-actin as protein loading control, protein sizes in kDa. (B) Quantification of peptide pre-adsorption indicates both full length mouse PANX1 (mPANX1-FL) and mPANX1-25K show significant competition with peptide (\**p*<0.05, \*\**p*<0.01). Two-way ANOVA followed by Šídák’s multiple comparisons test Bars represent mean ± SEM. *N*=3.

We wanted to confirm whether this isoform was also present in human cells and analyzed human embryonic kidney (HEK) and Hs578T breast cancer cell lines with and without CRISPR-Cas9 deletion of *hPANX1*. A 25 kDa immunoreactive species was detected in HEK and Hs578T cells regardless of the presence of hPANX1-FL when blots were probed with the PANX1 CT-412 antibody. However, unlike with mPANX-25K or hPANX1-FL, the band could not be competed with the cognate peptide (Figure 5A). The 25 kDa species was not seen when blots were probed with an NT PANX1 antibody, nor was the 25 kDa immunoreactive species detected in Western blots of GBM17, SCC-13, HEK *hPANX1* KO and Hs578T *hPANX1* KO human cell lines with the Sigma PANX1 CT antibody (Figure 5B). Taken together, this suggests the 25 kDa species is a mPANX1 isoform only.

**Figure 5.**
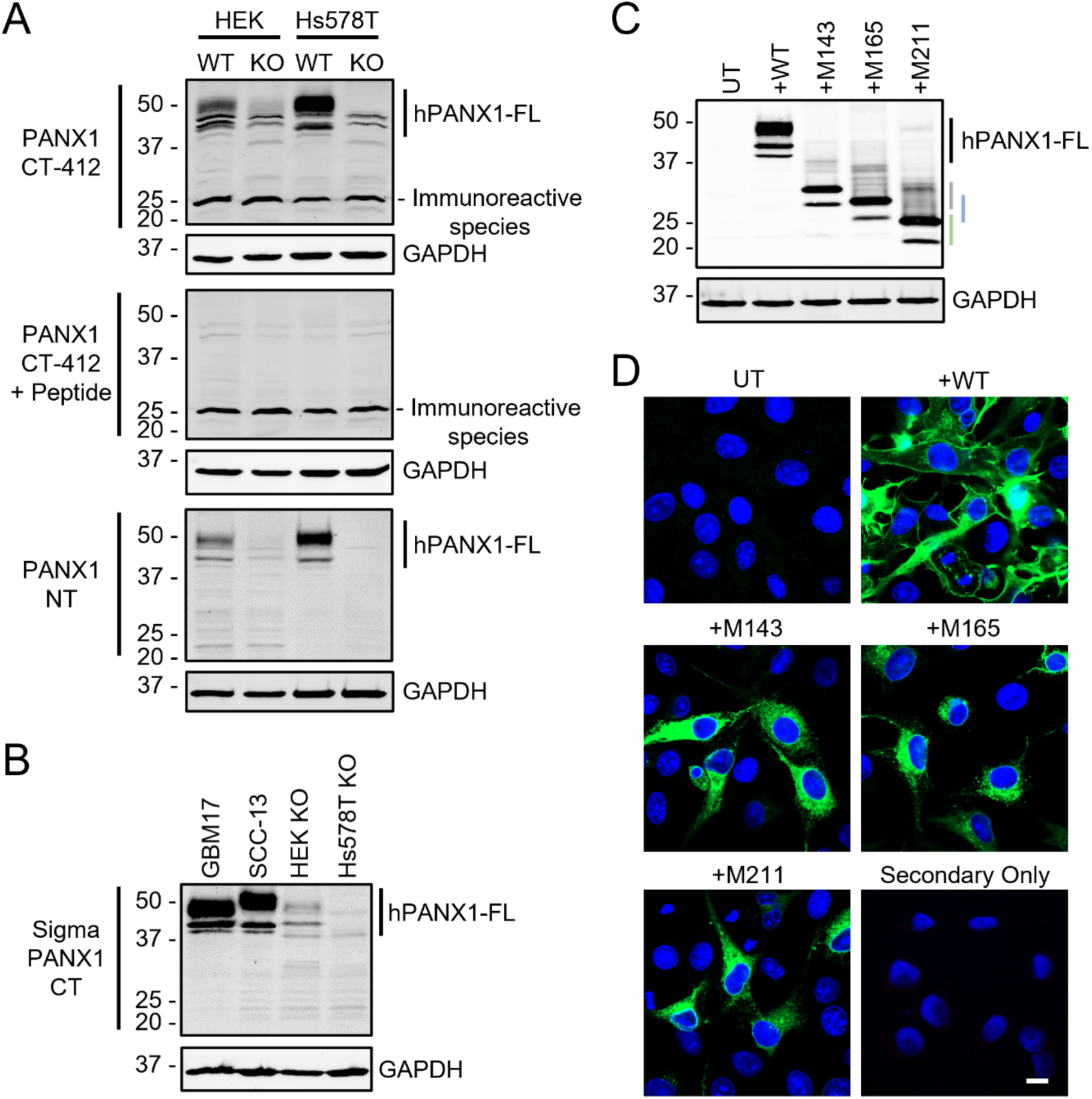
Human cell lines do not contain an endogenous 25 kDa species. (A) Immunoblotting of wildtype (WT) and full-length *hPANX1* knockout (KO) HEK293T and Hs578T cells. Blots probed with PANX1 C-terminal (CT)-412 antibody with and without peptide competition as well as a PANX1 N-terminal (NT) antibody. (B) Western blot of GBM17, SCC-13, HEK *hPANX1* KO and Hs587T *hPANX1* KO human cell lines with the Sigma PANX1 CT antibody. (C) Western blotting using the PANX1-CT-412 antibody shows Hs578T full-length *hPANX1* KO cells transfected with wildtype hPANX1 (WT), the shorter HA-tagged M143 (grey line), M165 (blue line) or M211 (green line) hPANX1 isoforms compared to untransfected (UT) controls. GAPDH as protein loading control. Protein sizes in kDa. Full-length human PANX1, hPANX1-FL. (D) Merged immunofluorescence microscopy images of hPANX1 isoforms in Hs578T full-length *hPANX1* KO cells. Coverslips probed with PANX1 CT-412 (green), Hoechst 33342 (nuclei in blue). Scale bar represents 10 µm. *N*=3.

We further validated these findings via transcriptomic analysis using Cap Analysis of Gene Expression sequencing (CAGE-seq) data (Figure 6A,B) from the FANTOM 5 project (Kanamori-Katayama *et al*., 2011; Consortium *et al*., 2014; Lizio *et al*., 2015) and Nanopore long-read RNA sequencing (Figure 6C,D) mined from a previous study (Weber *et al*., 2023). We found no evidence of alternative transcript usage or transcript start sites, suggesting that any potential mPANX1 or hPANX1 ATI species do not originate from an alternative transcription initiation site, alternative promoter usage or alternative transcript splicing of either *mPANX1* or *hPANX1*. This strengthens the possibility that mPANX1-25K arises from internal translation of the canonical *mPANX1* transcript instead of alternative transcription.

**Figure 6.**
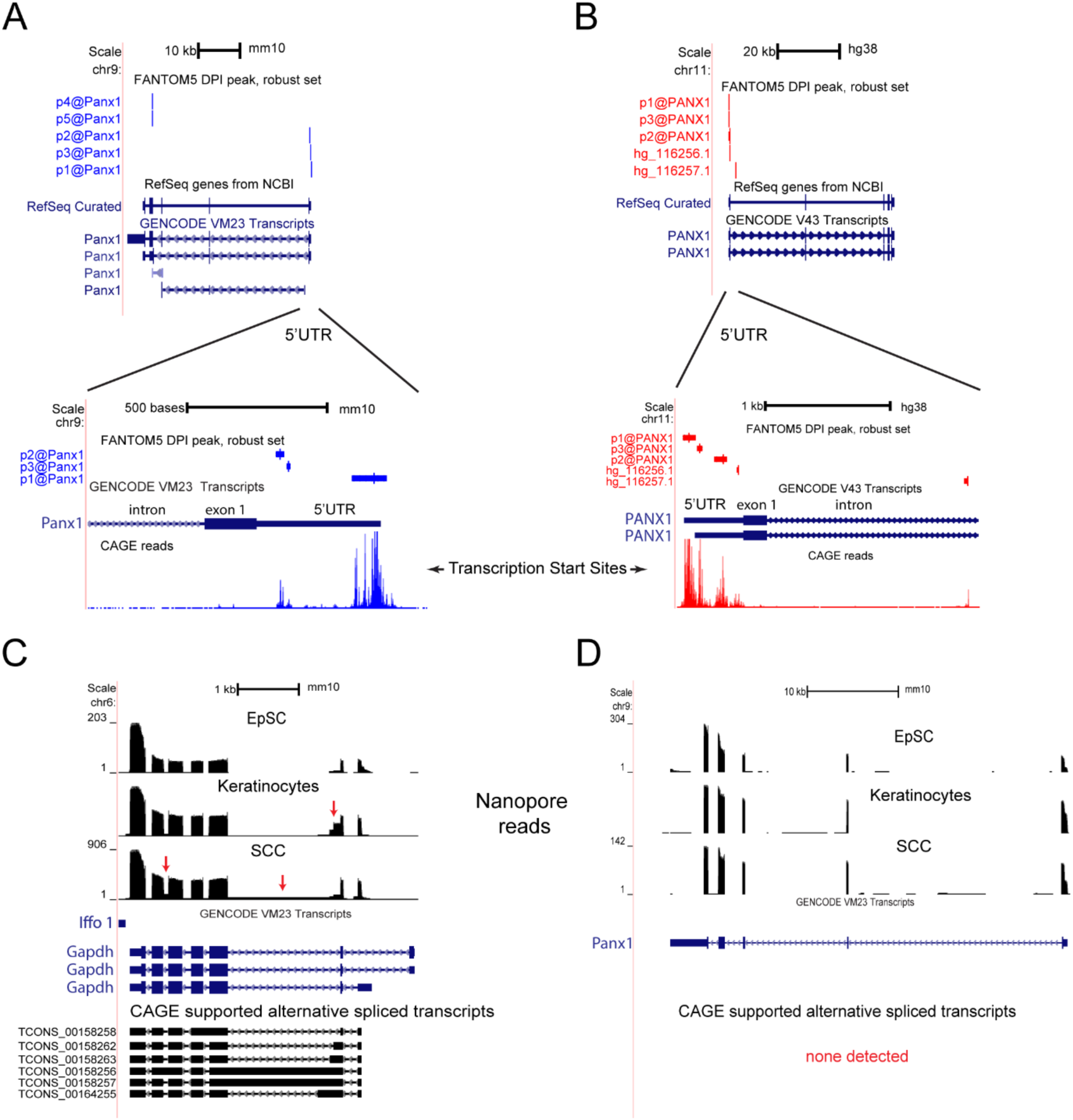
Transcriptome analysis suggests mPANX1-25K does not originate from alternate promoter usage or alternative splicing. (A) Cap Analysis of Gene Expression (CAGE-seq) peaks identify transcription start sites corresponding to promoter regions in the *mPanx1* 5’ untranslated region (UTR) and exon 4 of the canonical transcript (top). Predicted molecular weight of protein products resulting in transcripts originating in exon 4 are ∼7 kDa. Magnified view of alternate promoters within the 5’ UTR and intronic transcription start site peak with total CAGE-seq transcription start sites displayed (bottom). (B) CAGE-seq peaks identify transcription start sites corresponding to promoter regions in the *hPANX1* 5’ UTR, with an additional intronic peak near the 5’ end (top). Magnified view of alternate promoters within the 5’ UTR and intronic transcription start site peak with total CAGE-seq transcription start sites displayed (bottom). P@ = transcription start site clusters corresponding to promoter regions. (C) Nanopore long-read RNA sequencing identifies multiple alternative splicing events (red arrows) for *Gapdh* in epidermal stem cells (EpSCs), wildtype keratinocytes and cutaneous squamous cell carcinoma (SCC). Capped transcripts verified by combining CAGE-seq (CAGE supported alternative spliced transcripts). *Gapdh* was used as a positive control for alternative splicing events. (D) No CAGE-seq supported alternative spliced transcripts were identified for *mPanx1* using Nanopore long-read RNA sequencing in EpSCs, keratinocytes or SCCs.

### hPANX1 M211 exhibits intracellular localization, high mannose glycosylation and can interact with full-length hPANX1

With our discovery of the presence of a 25 kDa PANX1 isoform in mice but not human cells, as well as the sequence difference between mPANX1 M210 and hPANX1 I211, we wanted to further characterize potential mPANX1 and hPANX1 ATI isoforms. We designed hPANX1 mutant plasmids to investigate hPANX1 ATI variant trafficking, including constructs that contain only the hPANX1 sequence starting from M143 and M165. We also generated a construct containing the hPANX1 sequence starting at amino acid 211, but with a methionine instead of an isoleucine start (M211) to match the predicted 25 kDa isoform alternative start site mPANX1 M210 and force translation in an ectopic expression setting. WT, M143, M165, I211 and M211 constructs were expressed alone or M211 was co-expressed with WT in Hs578T *hPANX1* KO cells. While M143 (top species 32.9 kDa, lower species 30.3 kDa) and M165 (top species 30.91 kDa, lower species 28.02 kDa) were higher in molecular weight than endogenous mPANX1-25K, we found that M211 expression resulted in a banding pattern corresponding to mPANX1-25K (taking into consideration that the HA tag adds approximately 1 kDa to the molecular weight) with a doublet containing 26.0 kDa and 22.4 kDa species (Figure 5C). Through immunofluorescence microscopy analysis, we observed that hPANX1 WT exhibited prominent cell surface localization with some intracellular staining, whereas we identified a predominantly intracellular localization for M143, M165 and M211 (Figure 5D). Although I211 was barely detectable on Western blots containing other ectopically expressing cells due to adjustments made by imaging software and the presence of the 25 kDa immunoreactive species (Figure 7A), I211 levels were significantly increased compared to untransfected (UT) cells and appeared as a single species rather than a doublet, demonstrating that the isoleucine start site is capable of translation in this plasmid context, but less efficiently (Figure 7B). Through immunofluorescence microscopy analysis (Figure 7C), the I211 construct exhibited low transfection efficiency and faint intracellular expression. Thus, due to the increased expression compared to I211, we used the M211 plasmid for subsequent experiments to match the mPANX1 M210 start site. Furthermore, we observed hPANX1 WT exhibited plasma membrane localization with some intracellular staining, and we identified a largely cytoplasmic localization pattern for M211 as in Figure 5D, whereas in the WT and M211 co-expression condition, staining was observed at both the plasma membrane and intracellularly.

**Figure 7.**
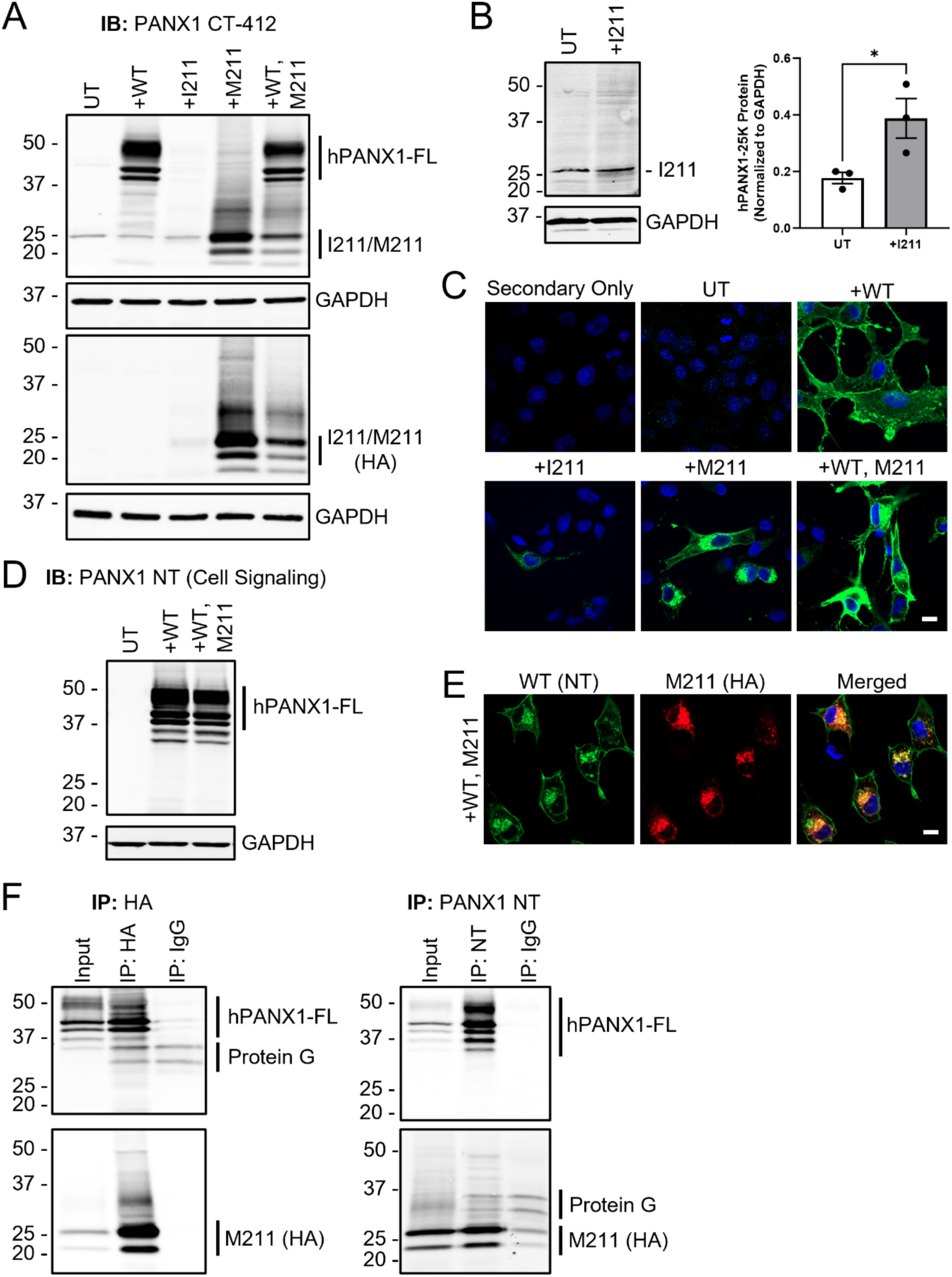
Overexpression of M211 demonstrates intracellular localization and association with hPANX1. (A) Western blotting using PANX1 CT-412 (top) and HA (bottom) antibodies shows Hs578T full-length *hPANX1* KO cells transfected with wildtype hPANX1 (WT), the shorter HA-tagged I211 or M211 hPANX1 isoforms or co-expression of the WT and M211 constructs compared to untransfected (UT) controls. (B) Immunoblotting at increased scanning intensity indicates I211 is significantly increased (\**p*<0.05) with I211 transient transfection compared to UT Hs578T full-length *hPANX1* KO cells. Unpaired t-test. Bars indicate mean ± SEM. (C) Merged immunofluorescence images of the hPANX1 isoforms in Hs578T full length *hPANX1* KO cells. Coverslips probed with PANX1 CT-412 (green). (D) Immunoblotting demonstrates hPANX1 WT but not M211 is detectable by PANX1 NT antibodies. GAPDH as loading control, protein sizes in kDa. (E) Confocal micrographs show hPANX1 WT (green, PANX1 NT) and M211 (red, HA) co-localization (yellow) in WT and M211 co-expressing Hs578T full-length *hPANX1* KO cells. (F) hPANX1 and M211 interaction shown by co-immunoprecipitation in WT, M211 expressing Hs578T full-length *hPANX1* KO cells using PANX1 NT (hPANX1) and HA (M211) antibodies. Rabbit and mouse immunoglobulin G (IgG) pulldown used as negative controls. Unspecific bands due to IgG antibodies in control and Protein G beads. Full-length human PANX1, hPANX1-FL. *N*=3.

To be able to assess for any co-localization between hPANX1-FL and M211, we used a PANX1 NT antibody which can detect hPANX1-FL but not M211 and an HA antibody which can only detect the HA-tagged M211 (Figure 7D). Similar to the PANX1 CT antibody immunofluorescence, we found hPANX1 WT exhibited both cell surface and intracellular localization while M211 appeared to be solely intracellular, with co-localization of WT and M211 in what appeared to be a Golgi cap structure (Figure 7E). Co-immunoprecipitation of hPANX1 WT and M211 confirmed this interaction in Hs578T *hPANX1* KO cells co-expressing the WT and M211 constructs (Figure 7F), indicating hPANX1-FL and M211 can associate. Interestingly, two smaller ∼30-37 kDa immunoreactive species containing the PANX1 NT epitope (Figure 7D) were also present in the co-immunoprecipitations and may correspond to M143 bands based on the predicted molecular weight (Figure 1). To further characterize the intracellular localization of M211, cells were co-stained with a variety of organelle markers including protein disulfide isomerase (PDI, endoplasmic reticulum lumen), calnexin (endoplasmic reticulum membrane), Golgi matrix protein 130 (GM130, cis/medial Golgi network), trans Golgi network 46 (TGN46), Ras-related protein 7 (RAB7, late endosome) or lysosomal-associated membrane protein 1 (LAMP1) in addition to an HA or PANX1 NT antibody (Figure 8). We observed a moderate degree of co-localization between M211 and calnexin, GM130 and TGN46, as well as hPANX1 WT and TGN46, indicating the interaction between hPANX1 WT and M211 most likely occurs in the endoplasmic reticulum and Golgi apparatus. M211 also showed strong co-localization with LAMP1 suggesting M211 is also present in lysosomes and may be degraded through this pathway which is common with construct overexpression.

**Figure 8.**
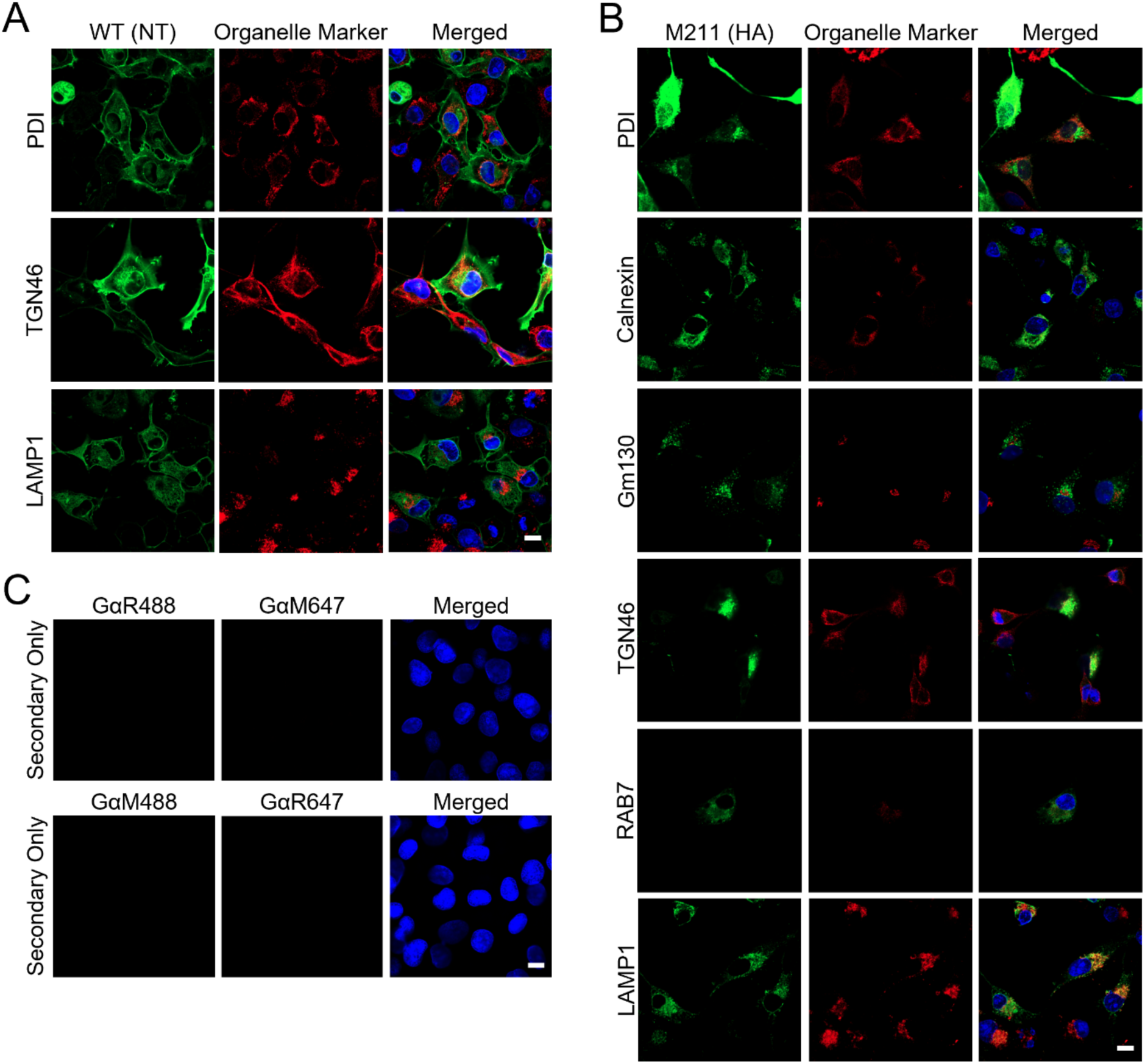
hPANX1 WT and M211 co-localization with various organelle markers. (A) Confocal micrographs of WT, M211 co-expressing Hs578T *hPANX1* KO cells show co-localization of PANX1 WT (PANX1 NT, A) or M211 (HA, B) in green with organelle markers for the endoplasmic reticulum (PDI, calnexin), Golgi apparatus (TGN46, Gm130), late endosome (RAB7) or lysosome (LAMP1) in red. Nuclei (blue) counterstained with Hoechst 33342. (C) Immunofluorescence images of coverslips incubated with Alexa Fluor™ 488 goat anti-rabbit, Alexa Fluor™ 488 goat anti-mouse, Alexa Fluor™ 647 goat anti-rabbit or Alexa Fluor™ 647 goat anti-mouse secondary antibody only as a secondary only control. Scale bars represent 10 µm. *N*=3.

As our findings suggest that M211 exhibits a predominantly intracellular localization, we next determined whether any proportion of M211 traffics to the plasma membrane. Cell surface markers wheat germ agglutinin (WGA) and E-cadherin were used to outline the plasma membrane in M211 or WT and M211-expressing Hs578T *hPANX1* KO cells (Figure 9A,B), and M211-expressing SCC-13 cells (Figure 9C,D). Cell surface line scans revealed that although not commonly seen, a subpopulation of M211 can reach the plasma membrane. Cell surface biotinylation in SCC-13 cells was consistent with the immunofluorescence analysis, demonstrating that a small subpopulation of M211 is present at the cells surface when M211 is overexpressed (Figure 9E).

**Figure 9.**
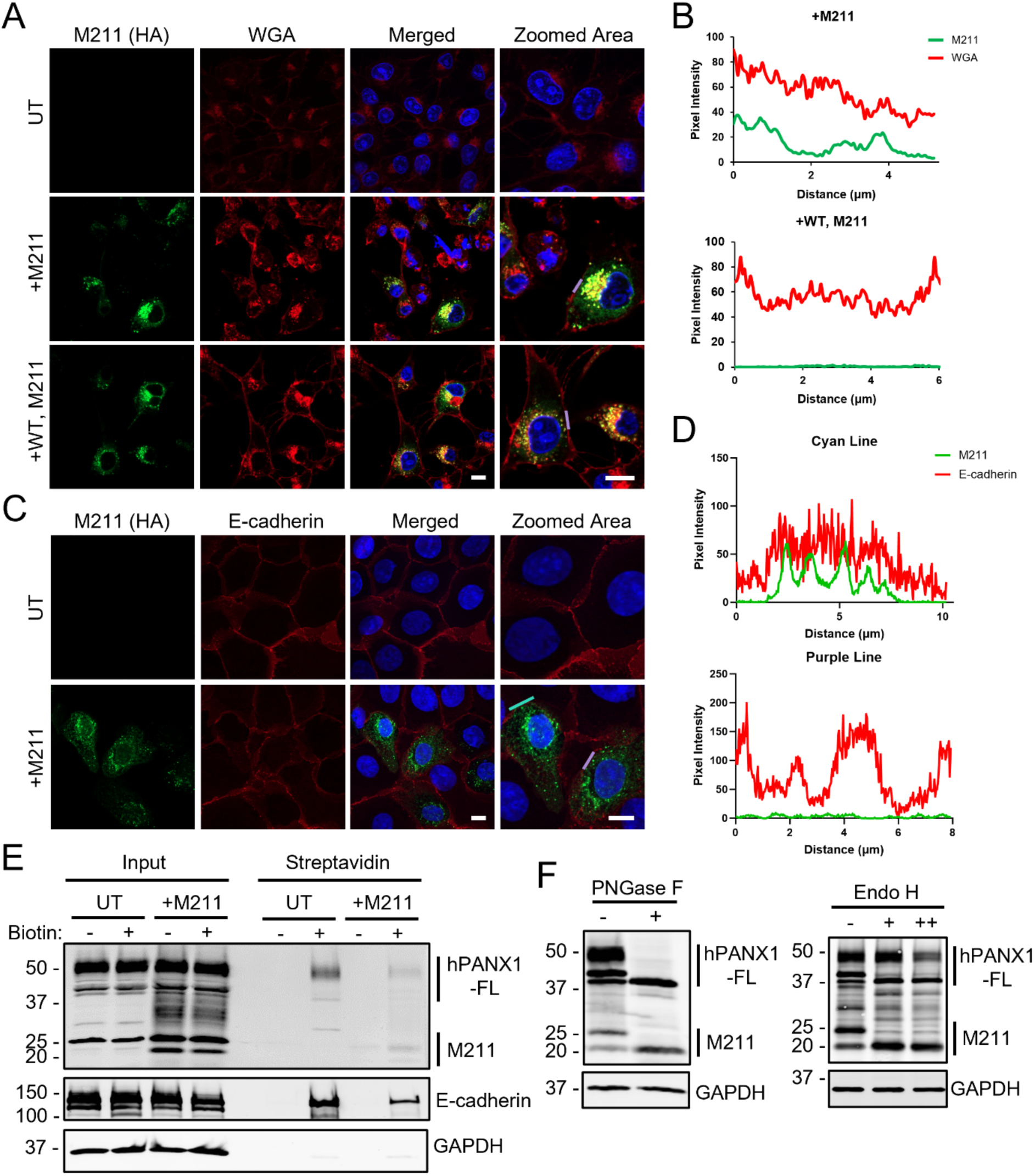
M211 is glycosylated and can traffic to the cell surface but is predominantly found in the cytoplasm. (A) Immunofluorescence images of untransfected (UT), M211 and wildtype (WT), M211 co-expressing Hs578T full-length *hPANX1* KO cells indicate M211 (HA, green) and wheat germ agglutinin (WGA, red) localization. Purple lines indicate areas for plasma membrane line scans. (B) Purple line scan quantifications identify some overlap of M211 and WGA staining at the cell surface in M211 only cells not seen with WT, M211 co-expression. (C) Confocal micrographs of UT or M211-expressing SCC-13 cells show localization of M211 (HA, green) and cell surface marker E-cadherin (red). Cyan and purple lines drawn at the plasma membrane mark areas for line scans. (D) Line scan quantifications indicate some overlap between M211 and E-cadherin at the plasma membrane. Scale bars represent 10 µm. Nuclei counterstained in blue with Hoechst 33342. (E) Cell surface biotinylation of UT and M211-expressing SCC-13 cells. E-cadherin used as positive control for cell surface localization, GAPDH as negative control (intracellular protein). (F) Both hPANX1-FL and M211 in Hs578T *hPANX1* KO cells co-expressing WT, M211 constructs are sensitive to PNGase F treatment via Western blot. Endo H digestion of Hs578T *hPANX1* KO cells co-expressing WT, M211 indicates partial digestion of M211 even when the amount of enzyme was doubled (++). GAPDH as loading control, protein sizes in kDa. *N*=3.

Since glycosylation can influence PANX1 trafficking (Penuela *et al*., 2009) and M211 retains the N255 glycosylation site of hPANX1-FL, we sought out to determine if M211 is glycosylated. To assess N-linked glycosylation (Figure 9F), we first treated lysates from Hs578T *hPANX1* KO cells co-expressing hPANX1 WT and M211 with PNGase F which removes all N-linked glycans. We saw a complete shift in all hPANX1 and M211 bands to the lowest molecular weight species indicating that M211 is glycosylated when ectopically expressed. To further characterize the glycan content, we treated the same lysates with Endoglycosidase H (Endo H) which cleaves only N-linked high mannose glycosylation. M211 exhibited only a partial digestion into the de-glycosylated form, even when the amount of enzyme was doubled, suggesting a small portion of M211 receives complex glycosylation but most of the species contain only high mannose glycans.

### mPANX1 ATI variants localize to the plasma membrane, but only M37 forms functional channels which require activation

To determine whether the alternatively translated mPANX1 proteins were also capable of trafficking to the plasma membrane and form channels on their own, we performed cell surface biotinylation assays (Figure 10A) and found a similar level of cell surface expression for mPANX1 translated using M1-only as compared with wildtype (WT) mPANX1. While the other mPANX1 variants could also be detected at the cell surface (M37-only, M143-only, and M210-only), the levels of the alternatively translated mPANX1 forms were relatively lower than the WT or M1-only constructs in total cell lysates and at the cell surface (Figure 10A), similar to hPANX1 M211 (Figure 9E). To determine whether different mPANX1 ATI forms can act as a functional ion channel at the plasma membrane, we performed whole-cell voltage clamp recordings in HEK293T cells expressing the mutant forms of mPANX1 (Figure 10B). We found that, similar to the WT mPANX1 (Sandilos *et al*., 2012), the M1-only form of mPANX1 was basally active in HEK293T cells, as evidenced by the presence of carbenoxolone (CBX)-sensitive, outwardly-rectifying currents. While both M1 and M37 translational initiation sites can produce a polypeptide with four intact transmembrane domains and are presumably capable of forming a complete mPANX1 channel complex, to our surprise, we did not observe any basal currents after expressing the M37-only form of mPANX1 in HEK293T cells. Similarly, the M143-only and M210-only mPANX1 forms did not display any channel activity. Notably, the whole-cell current of the M1-only form of mPANX1 displayed a more striking outwardly rectifying current-voltage (I-V) relationship than WT mPANX1, with no observable CBX-sensitive current at negative membrane potentials (Figure 10C). The mechanism for the altered I-V relationship remains unclear but is likely due to mutational effects of the downstream methionine substitutions.

**Figure 10.**
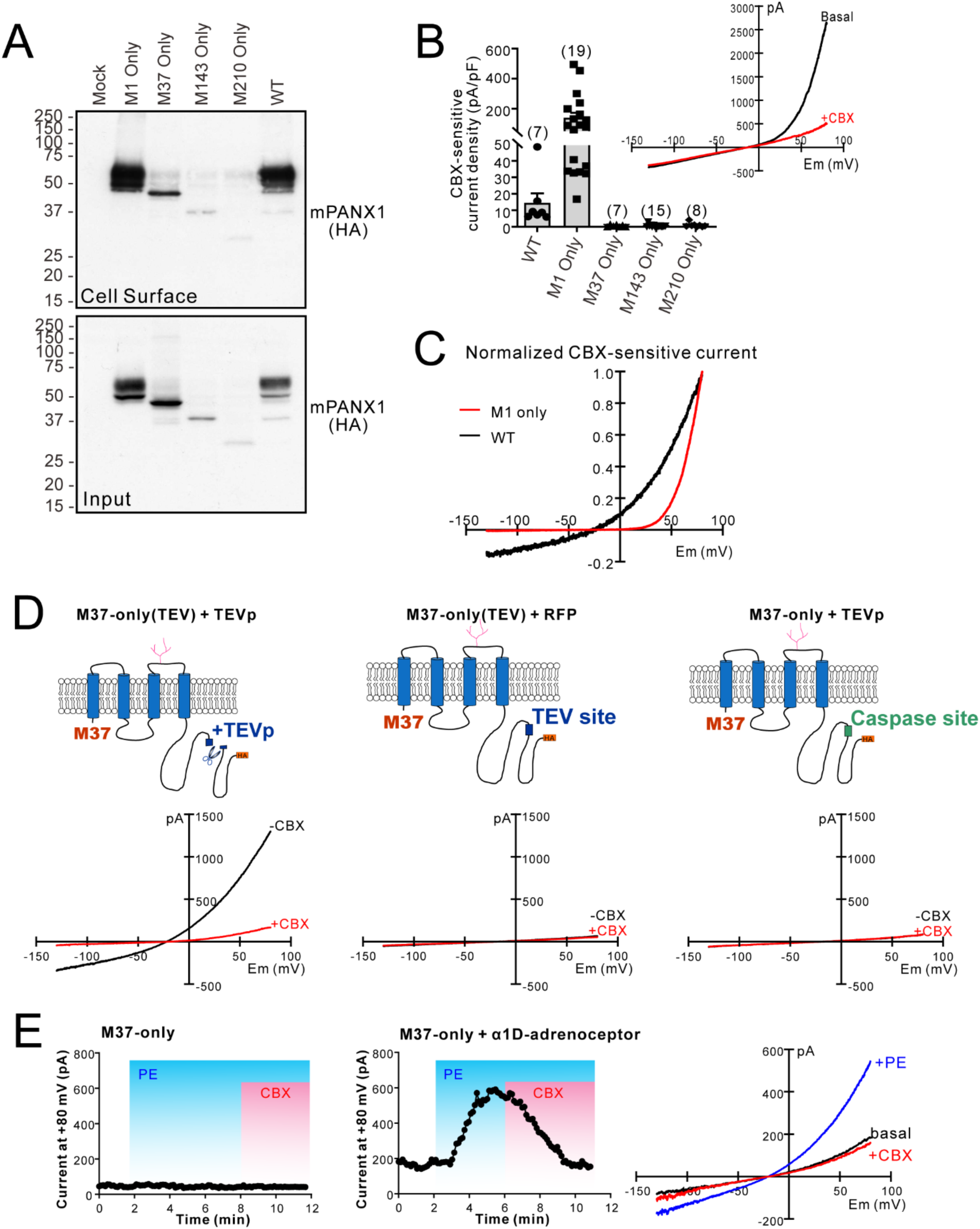
mPANX1 internal translation isoforms traffic to the cell surface, but only the M37-only channels are activated by C-tail cleavage or α1-adrenoceptor stimulation. (A) Input and cell surface biotinylation Western blot shows HEK293T cells transfected with wildtype mPANX1-HA (WT), shorter mPANX1 isoforms from each internal start site retaining only one in-frame methionine (M1, M37, M143, M210 Only) or empty vector (Mock) constructs. Tubulin used as protein loading control, protein sizes in kDa. (B) Whole-cell recordings of HEK293T cells transiently transfected with each mPANX1 isoform as indicated. Bracketed numbers show cell numbers recorded in each group. Inset: an exemplar whole-cell current of M1-only mPANX1, before (basal) and after bath application of 50 μM carbenoxolone (CBX). (C) CBX-sensitive whole-cell currents of WT or M1-only mPANX1 were normalized to its peak current at +80 mV. (D) Exemplar current-voltage (I-V) relationships of whole-cell recordings carried out in HEK293T cells co-expressing red fluorescent proteins (RFP) with either M37-only(Tobacco Etch Virus, TEV) and TEV protease (TEVp, left), M37-only(TEV) (middle), or M37-only and TEVp (right). Currents before (black line) and after the application of 50 μM CBX (red line) are shown. (E) Time series of exemplar whole-cell recordings of HEK293T cells expressing the M37-only form of mPANX1, with (middle) or without (left) co-expression of mouse α1D adrenoceptors. The I-V relationship of whole-cell currents (right) was obtained from a HEK293T cell co-expressing the M37-only form of mPANX1 and mouse α1D adrenoceptors under basal conditions (black line), after bath application of 20 μM phenylephrine (+PE; blue line) or after addition of 50 μM CBX (+CBX, red line).

To determine whether mPANX1 M37-only is capable of forming functional channels that require activation such as through the removal of the CT tail previously demonstrated for WT mPANX1 (Chekeni *et al*., 2011; Sandilos *et al*., 2012), we generated an M37-only(Tobacco Etch Virus, TEV) construct by replacing the CT caspase 3/7 cleavage site of mPANX1 M37-only with the TEV protease (TEVp) recognition sequence (Sandilos *et al*., 2012). Co-expression of M37-only(TEV) together with TEVp in HEK293T cells resulted in CBX-sensitive currents that were absent in cells co-expressing M37-only(TEV) and red fluorescence protein (RFP) or the M37-only construct with TEVp (Figure 10D). This indicates that mPANX1 M37-only can constitute functional cleavage-activated channels that are basally silent. To determine whether this form of mPANX1 can also be activated by receptor signaling pathways (Billaud *et al*., 2015), we co-expressed M37-only and the mouse α1D adrenoceptor in HEK293T cells and found that bath application of the α1 adrenoceptor agonist phenylephrine induced a CBX-sensitive PANX1 current that was not observed in cells expressing mPANX1 M37-only alone (Figure 10E). This further verifies that mPANX1 M37-only can form functional channels that require activation by multiple mechanisms. Unlike the strongly outwardly rectifying I-V relationship observed in cells expressing mPANX1 M1-only (Figure 10B inset), the I-V relationships observed from the TEVp-activated M37-only(TEV) channels (Figure 10D, left panel) and α1D adrenoceptor-activated M37-only channels (Figure 10E, right panel) were similar to those observed from mPANX1 WT (Sandilos *et al*., 2012; Billaud *et al*., 2015). Taken together, these findings illustrate that mPANX1 ATI isoforms mimic mPANX1-FL biochemistry, trafficking and/or channel function and some may exist to interact or regulate the full-length protein.

## DISCUSSION

PANX1 transcript isoforms have been documented in rodents and humans (O’Donnell and Penuela, 2023), and in breast cancer, the truncated PANX1^1-89^ mutant confers a metastatic advantage through augmented ATP release (Furlow *et al*., 2015; Wang *et al*., 2025). Many ATI isoforms of connexins have been identified and studied in various cell and disease contexts, but it was unknown whether pannexin ATI isoforms existed. Here, we report the initial characterization of multiple potential mPANX1 ATI isoforms, showing they are glycosylated and can traffic to the cell surface, but not all can make functional channels. Furthermore, we identified a 25 kDa mPANX1 isoform which does not have an endogenous hPANX1 equivalent, but when the analogous hPANX1 M211 is expressed in human cells it exhibits an intracellular localization, can be N-linked glycosylated and interacts with hPANX1-FL.

Our results suggest that the *mPANX1* ATG sequence corresponding to M210 is likely the mPANX1-25K start site since mPANX1-25K matches the predicted molecular weight of the M210 protein. mPANX1-25K could arise from alternative translation of the full-length *mPANX1* transcript since we could not identify any alternate promoter start sites or transcript variants which would account for a mPANX1 isoform of that size in our own analysis. However, we recognize that the transcriptomic analysis may not be representative of the cell lines used in our own study, and the above results do not preclude mPANX1-25K from being produced from alternative transcription, splicing or promotor usage, meaning we cannot rule out the possibility of a transcript or splice variant in other cells. Furthermore, mPANX1-25K is detected by antibodies targeting PANX1 CT but not NT. To further investigate the possibility of mPANX1-25K as an ATI mPANX1 isoform beginning at M210, future studies should perform ribosome sequencing of the *mPanx1* transcript to positively identify internal translation start sites present throughout the mRNA. Furthermore, there is evidence of a predicted rat *Panx1* transcript variant X1 (XM_039081351.2) which could produce the predicted rat PANX1 isoform X1 (XP_038937279.1) that includes rat PANX1 residues M210-C426 (O’Leary *et al*., 2016). This variant has not been confirmed at the protein level, but the sequence spans the exact residues we predict for mPANX1-25K. Deep RNA sequencing would be needed to identify a potential mPANX1-25K transcript, especially if the transcript is lowly expressed (Goldman and Domschke, 2014). Lastly, mPANX1 contains only one caspase cleavage sites at aspartic acid residue D378 (Sandilos *et al*., 2012) which would produce CT fragments detectable by the PANX1 CT-412 antibody used in our study. However, the cleavage product would not correspond to the ∼25 kDa molecular weight of the mPANX1-25K isoform since proteolysis at D379 would produce a much shorter product (∼5.2 kDa).

We initially included hPANX1 I211 as a potential internal start site in our analysis since it corresponds to a methionine in mice (M210), but recognize it constitutes an alternative non-AUG start. Although there are not many reports of mammalian proteins starting with isoleucine, one well-known example is the transcriptional enhancer factor TEF-1 (also known as TEA domain family member 1, TEAD1) which is endogenously translated beginning at an AUU start codon which also encodes isoleucine (Xiao *et al*., 1991). New evidence suggests that translation initiation events at near-cognate codons are much more common than initially believed, accounting for up to 50% of initiation, even though translation from non-AUG codons such as AUC is generally less efficient (Kearse and Wilusz, 2017; Diaz de Arce *et al*., 2018). However, we were unable to identify an endogenous 25 kDa hPANX1 species corresponding to an I211 start site that could be competed with peptide pre-adsorption. This is consistent with the finding that the *hPANX1* mRNA sequences flanking the I211 AUC start site do not completely match the Kozak consensus sequence (Kozak, 1986) which can have considerable influence for translation initiation efficiency for non-AUG codons (Kearse and Wilusz, 2017; Diaz de Arce *et al*., 2018). It should be noted that the mRNA sequence flanking the M210 AUG start site does not fully match the Kozak consensus sequence either (Kozak, 1986), but there may be other factors to facilitate this internal translation. Factors such as internal ribosome entry sites, cap-independent translation elements or mRNA structures adjacent to the start site are common mechanisms used by cells to promote ATI (James and Smyth, 2018), but would need to be investigated further in the case of *mPANX1* mRNA. Interestingly, although all primates contained an isoleucine at residue 211, there was an approximately even split between the rodent species we investigated as to whether the codon for methionine or valine was present at this site. Valine codons, particularly GUG which differs from the canonical AUG by only one base pair, have also been shown to act as a translation site in eukaryotic organisms. For example, eukaryotic translation initiation factor 4G2, which functions in ATI via internal ribosome entry sites, solely uses a GUG start codon for translation initiation (Kearse and Wilusz, 2017). In another study looking at initiator tRNAs, it was found that tRNAs loaded with valine (containing the anticodon GAC which corresponds to the GUC mRNA codon) can function efficiently as an initiator tRNA and is most likely recruited by eukaryotic translation initiation factor 2D (Drabkin and RajBhandary, 1998; Kearse and Wilusz, 2017). Thus, even though the *mPANX1* sequence diverged in the rodent family, it is possible that the 25 kDa PANX1 species may still be present in the species beginning with the non-AUG valine start site.

There are many possibilities for the function of mPANX1-25K as a novel isoform in mouse cells. M211 localizes predominantly to the Golgi network where it interacts with hPANX1-FL, rarely trafficking to the cell surface. This is similar to GJA1-20k, which localizes to the endoplasmic reticulum and Golgi apparatus where it associates with Cx43 but does not reach the cell surface to be incorporated into gap junction plaques (Smyth and Shaw, 2013; James and Smyth, 2018). Thus, like GJA1-20K and Cx43, M211 could interact with hPANX1-FL in the Golgi network to autoregulate its oligomerization and trafficking within the cell. On the contrary, we have seen in melanoma cells that there is increased abundance of intracellular hPANX1 rather than at the plasma membrane (Freeman *et al*., 2019; Sayedyahossein *et al*., 2021), so it is possible that instead of aiding hPANX1-FL trafficking to the cell surface, M211 binds to hPANX1-FL to sequester the protein intracellularly. Moreover, since the protein may contain multiple transmembrane domains, it is possible that mPANX1-25K could oligomerize to form a channel. However, based on the electrophysiology data obtained with the mouse expression constructs, we would expect that only M1 and M37 are able to form functional channels on their own. It is more likely that the mPANX1-25K isoform could intermix with full length PANX1 to change the structure and activity of the mixed channel similar to the PANX1^1-89^ mutant (Furlow *et al*., 2015), although a homomeric PANX1^1-89^ channel has been shown to be functional (Wang *et al*., 2025). Since we saw that hPANX1-FL and M211 interact by co-immunoprecipitation, it is possible that mPANX1-25K could associate with mPANX1-FL. Future studies should investigate which domains of mPANX1-FL and mPANX1-25K interact and the potential relative stoichiometry of each mPANX1 form in a mixed channel. Hypothetically, a channel containing mPANX1-25K could be hyperactive since this isoform lacks the NT domain thought to be critical for channel gating (Mou *et al*., 2020), which would be advantageous to cancer cells, for example, since channel inhibition via PANX1 channel blockers CBX, probenecid and spironolactone reduce cancer cell properties such as growth, migration, invasion and epithelial to mesenchymal transition in multiple cancer types (Freeman *et al*., 2019; Jalaleddine *et al*., 2019; Sun *et al*., 2020). However, the proposed NT-mediated gating mechanisms, either through obstructing the small-ion-permeating side tunnels (Ruan *et al*., 2020) or rearranging NT conformation within the PANX1 permeation pathway (Kuzuya *et al*., 2022) should be revisited. Given our findings that the NT-omitted M37-only form of mPANX1 can be still activated by reversible and irreversible mechanisms, and that M37-only(TEV) and TEVp co-expression resulted in whole-cell currents with a similar I-V curve to the CT-cleavage-activated full-length channels, the unidentified mPANX1 gating apparatus is likely found between M37 and M372 residues of PANX1. Lastly, through its interaction with the hPANX1-FL channel, M211 could either enhance or inhibit binding of protein interactors to hPANX1-FL. This could have signalling consequences in cells, especially in cancers like melanoma where hPANX1-FL functions in Wnt signalling through its direct interaction with β-catenin to drive melanoma cell metabolism and proliferation (Sayedyahossein *et al*., 2021).

Through enzymatic assays, we determined exogenously expressed M211 shows biochemical similarities to hPANX1-FL such as its ability to undergo N-linked glycosylation. Post-translational modification of mPANX1 such as glycosylation and phosphorylation have been shown to alter channel trafficking (Penuela *et al*., 2007; Penuela *et al*., 2009) and activity (Weilinger *et al*., 2012; Weilinger *et al*., 2016), respectively, which may also occur in the case of mPANX1-25K. To this end, Endo H and PNGase F digestion of M211 determined a large proportion of this isoform contains high mannose glycans, but a small fraction is further modified to complex glycosylation. This is supported by M211 being found predominantly in intracellular compartments like the Golgi network, corresponding to the high mannose Gly1 species of hPANX1-FL which is present in the endoplasmic reticulum and early Golgi apparatus. In the same way, a small fraction of M211 reaches the plasma membrane similar to hPANX1 Gly2 which is complexly glycosylated and predominantly found at the cell surface (Penuela *et al*., 2007; Penuela *et al*., 2009). Based on our prediction that endogenous mPANX1-25K originates from canonical *mPANX1* beginning at the M210 start site, mPANX1-25K would retain an identical protein sequence and corresponding modification and protein interaction sites present after the M210 residue. Future work should further characterize the biochemical properties of mPANX1-25K such as phosphorylation.

Our finding that a 25 kDa isoform is present for mPANX1 but not hPANX1 reiterates that there are some distinct species-specific differences for PANX1. The majority of early studies investigating PANX1 post-translational modifications used mPANX1 or rat PANX1, including the identification of the site of N-linked glycosylation at N254 (Penuela *et al*., 2007) and phosphorylation on residues Y198 (Lohman *et al*., 2015; DeLalio *et al*., 2019), S205 (Poornima *et al*., 2015), T302 (Lopez *et al*., 2020), Y308 (Weilinger *et al*., 2012; Weilinger *et al*., 2016), S328 (Lopez *et al*., 2020) and S394 (Poornima *et al*., 2015). Apart from a recent study investigating hPANX1 Y199 and Y309 phosphorylation—to which they believe SRC phosphorylation does not occur at these sites in hPANX1 (Ruan *et al*., 2024)—there have been no other studies to our knowledge that have confirmed the mPANX1/rat PANX1 post-translational modifications translate to hPANX1. Furthermore, mPANX1 contains only one caspase cleavage site at D378 (Penuela *et al*., 2014) whereas hPANX1 contains two caspase cleavage sites at aspartic acid residues D167 and D379 (Chekeni *et al*., 2011; Sandilos *et al*., 2012; Boyce *et al*., 2018). This means that even though mPANX1 and hPANX1 have 94% sequence conservation (Baranova *et al*., 2004b), we cannot assume that the same post-translational processing of PANX1 occurs in each species. Furthermore, there may be differences in the post-translational modifications present in different mPANX1 isoforms and further research should be conducted to investigate if mPANX1-25K undergoes the same PANX1 modifications as mPANX1-FL.

To our knowledge, this is the first report and characterization of the mPANX1-25K isoform that may act as a regulator of the mPANX1-FL channel. Our results emphasize the importance of investigating additional isoforms of PANX1 as well as being cautious to assume properties of mPANX1 translate to hPANX1, considering that isoforms like mPANX1-25K may change the landscape of what is known about PANX1 biochemistry, trafficking and function.

## MATERIALS AND METHODS

### PANX1 sequence analysis

NCBI RefSeq mRNA and protein sequences of mPANX1 (mRNA NM_019482.2, protein NP_062355.2) and hPANX1 (mRNA NM_015368.4, protein NP_056183.2) (O’Leary *et al*., 2016) were analyzed to identify potential internal methionine and isoleucine start sites downstream of the canonical M1 site. mRNA sequences of mPANX1 and hPANX1 were translated using Expasy (Duvaud *et al*., 2021) to visualize the protein sequence which allowed us to identify neighbouring amino acid residues and corresponding codons of predicted methionine and isoleucine start sites to compare to the Kozak consensus sequence (Kozak, 1986). Protein sequence homology was compared using NCBI standard protein BLAST, inputting hPANX1 protein as the query sequence. Aligned sequences were compared between hPANX1 and relevant organisms with available complete PANX1 sequences and included based on phylogenetic relevance and sequence homology to *Homo sapiens*. Theoretical molecular weights were calculated using the Expasy (Duvaud *et al*., 2021) Compute pI/Mw (isoelectric point/molecular weight) Tool (https://web.expasy.org/compute_pi/).

### Cell culture

HEK293T (ATCC CRL-3216, RRID:CVCL_0063), HEK293T *hPANX1* KO (Nouri-Nejad *et al*., 2021), Hs578T (ATCC HTB-126, RRID:CVCL_0332), Hs578T *hPANX1* KO (Nouri-Nejad *et al*., 2021) and GBM17 cell lines were cultured in DMEM media (Gibco #12430062) supplemented with 10% fetal bovine serum (FBS; Thermo Fisher Scientific #12483020) and 1% penicillin/streptomycin (Gibco #15140-122). B16-F0 (ATCC CRL-6322, RRID:CVCL_0604), B16-F10 (ATCC CRL-6475, RRID:CVCL_0159), and B16-BL6 (kindly provided by Dr. Moulay Alaoui-Jamali (McGill University), RRID:CVCL_0157) murine melanoma cell lines were cultured in DMEM containing 10% FBS, 100 units/ml penicillin and 100µg/ml streptomycin. SCC-13 cells (RRID:CVCL_4029; gifted by Dr. Lina Dagnino, University of Western Ontario) were cultured in DMEM media supplemented only with 10% FBS (WISENT #098-150). Cells were maintained at 37°C with 5% CO_2_ and confirmed mycoplasma-negative using the MycoAlert® Mycoplasma Detection Kit (Lonza #LT07-318).

### Cloning and plasmids

The mPANX1-HA expression vector was a gift from Dr. Kodi Ravichandran (Washington University) (Chekeni *et al*., 2011; Billaud *et al*., 2015). A different Kozak sequence was used in the construct encoding HA-tagged mPANX1 (Gcc) at the M1 site compared to the canonical *mPANX1* transcript (Acc). The Tobacco Etch Virus protease (TEVp) plasmid was kindly provided by S. R. Ikeda (Williams *et al*., 2009), and the pRFP-C1 plasmid was kindly provided by Dr. Julius Zhu (University of Virginia). The mouse α1D adrenoceptor plasmid was obtained from OriGene (#MR222643) (Chiu *et al*., 2017). To generate different methionine to isoleucine mutations, site-directed mutagenesis was performed on the mPANX1-HA construct using PfuTurbo polymerase (Agilent #600250) with various combinations of primers (Table 1). To generate the M37-only(TEV) constructs, the CT caspase 3/7 cleavage site of M37-only form of mPANX1 (IKMDIID) was replaced by TEVp recognition sequence (ENLYFQG) using the same site-directed mutagenesis approach as above. The hPANX1 WT plasmid was purchased from InvivoGen (#puno1-hpanx1) and modified by NorClone Biotech Laboratories to create the vectors encoding the M143, M165, I211 and M211 proteins which were tagged with HA on the CT. All constructs were verified by Sanger sequencing.

**Table 1.**
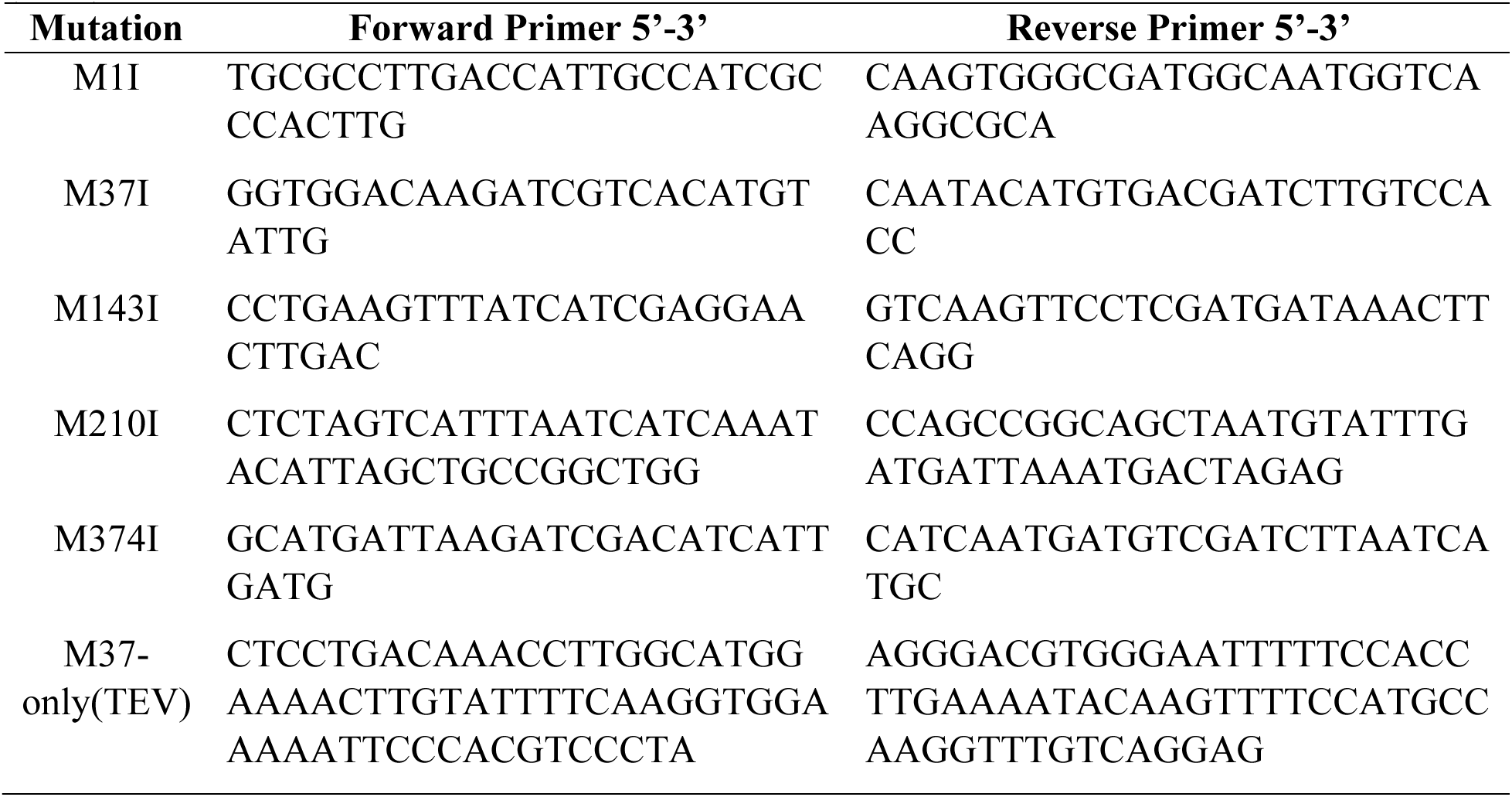
Primers used for generating methionine-to-isoleucine mutations and Tobacco Etch Virus (TEV) in mPANX1.

### Transfection

Transient transfections were performed using Lipofectamine™ 2000 (Invitrogen #11668027) for all mouse plasmids and Lipofectamine™ 3000 (Invitrogen #L3000015) for all human plasmids. For immunoblotting and cell surface biotinylation of mPANX1, 4 µg and 8 µg of mPANX1 plasmids were used for transfections in a 6-well plate or 6-cm dish, respectively. For whole-cell recordings, cells were co-transfected with 0.9 µg of a plasmid containing mouse adrenergic α1D receptor (OriGene #MR222643) (Chiu *et al*., 2021) or TEVp (Sandilos *et al*., 2012), 0.3 µg of mPANX1 constructs and 0.1 µg of EGFP-encoding plasmids (pEGFP-C1) in a 12-well plate. Unless otherwise stated, 0.5 µg of all hPANX1 plasmids (0.25 µg each of hPANX1 WT and M211 in the co-expression condition) were used for transfections in a 6-well plate. Reagents were added to Opti-MEM media (Life Technologies #51985-034) and cells were transfected according to the manufacturer’s protocol. Cells were transfected 24 h after seeding and experiments were performed 24 h or 48 h after transfection as noted.

### Protein isolation and immunoblotting

In HEK293T cells overexpressing various mPANX1 plasmids, protein was extracted using a PBS buffer containing 1% Triton X-100, protease inhibitor cocktail (Sigma-Aldrich #P8340), 10 mM NaF and 10 mM NaVO_3_. Unless otherwise noted, protein was isolated from human cell lines using 1x RIPA buffer (50 mM Tris-HCl at pH 8.0, 150 mM NaCl, 1% Nonidet™ P 40 Substitute [Sigma-Aldrich #74385], 0.5% sodium deoxycholate). Extraction buffers were supplemented with 1 mM NaF, 1 mM Na_3_VO_4_ and half of a Pierce Protease Inhibitor EDTA-free Mini Tablet (Thermo Fisher Scientific #A32955). Protein was quantified using a bicinchoninic acid assay (BCA; Thermo Fisher Scientific #23225).

For immunoblotting analyses of mPANX1 plasmids, 50 µg or 60 µg protein samples were separated by SDS-PAGE using an 8% acrylamide gel, transferred onto 0.45 µm nitrocellulose membranes (PerkinElmer, Inc. #NBA085B001EA) and the membranes were blocked with 5% non-fat dry milk dissolved in a tris-based buffer (10 mM Tris, 150 mM NaCl, and 0.1% Tween 20, pH 7.4) at room temperature for 1 h. HA-tagged mPANX1 isoforms were probed with anti-HA antibody 1:1000 (Cell Signaling Technology #2367, RRID:AB_10691311) and anti-α-tubulin antibody 1:8000 (Sigma-Aldrich #T9026, RRID:AB_477593) as a loading control. Amersham ECL horseradish peroxidase (HRP)-linked secondary antibodies (GE Healthcare #NA931V or #NA9340V, 1:4000–1:8000) and Western Lightning Plus-ECL (PerkinElmer, Inc. #NEL103001EA) were used to visualize immunoreactive signals on Amersham Hyperfilm ECL (GE Healthcare #28-9068-39). Blots were digitized using the Epson Perfection 4990 Photo desktop scanner. Primary antibodies were diluted in 0.1% Tween 20 PBS and incubated overnight at 4°C and secondary antibodies were dissolved in 0.1% Tween 20 PBS with 5% non-fat dry milk and incubated for 1 h at room temperature.

Western blots for hPANX1 experiments and endogenous mPANX1 experiments were performed as previously reported (O’Donnell *et al*., 2025a), loading 40-50 µg to analyze endogenous proteins from all cell types, except for B16-F0, F10 and BL6 cells where 70 µg was loaded per sample. For Hs578T *hPANX1* KO overexpressing cells, 40 µg per sample was used. Protein samples were separated on 10% acrylamide SDS-PAGE gels which were transferred to nitrocellulose membranes using an iBlot Gel Transfer Device (Invitrogen) on P3. After blocking for 1 h at room temperature in 1x PBS with 3% bovine serum albumin (BioShop ALB001.100), most antibodies were incubated overnight at 4°C, except GAPDH, β-actin and secondary antibodies which were incubated for 1 h at room temperature. Primary antibody working dilutions included: anti-PANX1 CT-395 1:1000, anti-PANX1 CT-412 1:1000 (Penuela *et al*., 2007; Penuela *et al*., 2009), anti-PANX1 NT 1:1000 (Cell Signaling Technology #91137, RRID:AB_2800167), anti-PANX1 CT 1:1000 (Sigma-Aldrich #HPA016930), mouse anti-HA 1:5000 (Invitrogen #26183, RRID:AB_2533049), rabbit anti-HA 1:1000 (Cell Signaling Technology #3724, RRID:AB_1549585), mouse anti-GAPDH 1:5000 (Millipore Sigma #G8795, RRID:AB_1078991), mouse anti-β-actin 1:5000 (Millipore Sigma #A5441, RRID:AB_476744) and mouse anti-E-cadherin 1:2500 (BD Transduction Laboratories™ #610182, RRID:AB_397580). Dilutions of 1:10,000 were used for all secondary antibodies, including IRDye-800CW goat anti-rabbit (LI-COR Biosciences #926-32211, RRID:AB_621843), IRDye-800CW goat anti-mouse (LI-COR Biosciences #926-32210, RRID:AB_621842), IRDye-680RD goat anti-mouse (LI-COR Biosciences #926-68070, RRID:AB_10956588) and IRDye-680RD goat anti-rabbit (LI-COR Biosciences #926-68071, RRID:AB_10956166) immunoglobulin G (IgG) secondary antibodies. All antibodies were dissolved in 3% bovine serum albumin in 0.05% Tween 20 PBS. Blot imaging and protein and molecular weight quantifications were performed using the LI-COR Odyssey Infrared Imaging System and ImageStudio software (LI-COR Biosciences). Protein levels were normalized to GAPDH and presented relative to the control mean value.

### Peptide pre-adsorption

Peptide pre-absorption was performed by incubating the diluted mouse PANX1 CT-375 or human PANX1 CT-412 antibody with 50:1 molar excess of the cognate mouse or human PANX1 peptide (Penuela *et al*., 2007; Freeman *et al*., 2019), respectively, for 30 min at room temperature before immunoblotting.

### Immunocytochemistry

For analysis of hPANX1 isoform localization in Hs578T *PANX1* KO cells, 250,000 cells were plated on glass coverslips (Fisher Scientific #1254580) in a 6-well plate. Forty-eight h after transfection, cells were fixed with ice-cold methanol:acetone (5:1, v/v) for 15 min at 4°C after two washes with 1x DPBS without Ca^2+^ and Mg^2+^. In a humidity chamber, coverslips were blocked with 10% goat serum (Life Technologies #50062Z) for 1 h at room temperature, and then incubated overnight at 4°C with anti-PANX1 NT 1:250 (Cell Signaling Technology #91137, RRID:AB_2800167), anti-PANX1 CT-412 1:500 (Penuela *et al*., 2007; Penuela *et al*., 2009), rabbit anti-HA 1:500 (Cell Signaling Technology #3724, RRID:AB_1549585) and/or mouse anti-HA 1:500 (Invitrogen #26183, RRID:AB_2533049). All antibodies were diluted in 1% goat serum in 1x DPBS without Ca^2+^ and Mg^2+^. The next day, coverslips were washed thrice for 5 min using 1x DPBS without Ca^2+^ and Mg^2+^, probed with 1:700 Alexa Fluor™ 488 goat anti-rabbit (Invitrogen #A-11008, RRID:AB_143165), 1:700 Alexa Fluor™ 488 goat anti-mouse (Invitrogen #A-11029, RRID:AB_2534088), 1:500 Alexa Fluor™ 647 goat anti-rabbit secondary antibody (Invitrogen A-21244, RRID:AB_2535812) and/or 1:500 Alexa Fluor™ 647 goat anti-mouse secondary antibody (Invitrogen #A-21236, RRID:AB_2535805) for 1 h at room temperature. After 5 min 1x DPBS without Ca^2+^ and Mg^2+^ and distilled water washes, coverslips were stained for cell nuclei with Hoechst 33342 (Life Technologies #H3570) diluted 1:1000 in double-distilled water for 5 min at room temperature. Coverslips were mounted with ProLong™ Gold Antifade Mountant (Thermo Fisher Scientific #P36934) and imaged on a ZEISS LSM 800 AiryScan Confocal Microscope (Schulich Imaging Core Facility, University of Western Ontario) with 63x oil magnification and 405 nm (Hoechst 33342), 488 nm (Alexa Fluor™ 488) and 640 nm (Alexa Fluor™ 633, 647) laser lines.

Mouse anti-E-cadherin 1:200 (BD Transduction Laboratories™ #610182, RRID:AB_397580), and Alexa Fluor™ 633-conjugated WGA 1:100 (Thermo Fisher Scientific #W21404) were incubated with coverslips for 1 h at room temperature to label the cell surface of SCC-13 and Hs578T *PANX1* KO transfected cells, respectively. In addition to labeling the cell surface, it should be noted that WGA also binds N-acetylglucosamine and N-acetylneuraminic acid residues present in the Golgi apparatus. Line intensity profiles of fluorescent images (line scans) were quantified using ImageJ (Fiji) (Schindelin *et al*., 2012) and visualized for overlap of pixel intensities from each channel. Mouse anti-GM130 1:150 (BD Transduction Laboratories #610822, RRID:AB_398141), mouse anti-TGN46 1:75 (Thermo Fisher Scientific #MA3-063, RRID:AB_325484), mouse anti-LAMP1 1:100 (Thermo Fisher Scientific #14-1079-80, RRID:AB_467426) were incubated on coverslips to visualize the cis/medial-Golgi network, trans Golgi network and lysosomes, respectively. Mouse anti-PDI 1:150 (Enzo Life Science #ADI-SPA-891F, RRID:AB_10615355) and rabbit anti-calnexin 1:100 (Cell Signaling Technology #2679S, RRID:AB_2228381) were used to visualize the lumen and membrane of the endoplasmic reticulum, respectively. Rabbit anti-RAB7 antibody 1:75 (Cell Signaling Technology #9367, RRID:AB_1904103) was used to visualize late endosomes. All organelle markers were incubated on coverslips for 1 h at room temperature as previously described (Leighton *et al*., 2024).

### Transcriptomics

The FANTOM 5 project (Kanamori-Katayama *et al*., 2011; Consortium *et al*., 2014; Lizio *et al*., 2015) reprocessed data (https://fantom.gsc.riken.jp/5/datafiles/reprocessed/) was used for the CAGE-seq analysis. Transcription start site (CAGE-seq) peaks across the panel of the biological states (samples) were identified by decomposition based peak identification (Consortium *et al*., 2014), where each of the peaks consists of neighbouring and related transcription start sites. Nanopore long-read RNA sequencing (Weber *et al*., 2023) of P60 mouse epidermal stem cells, keratinocytes, and HrasG12V; *Tgfbr2* KO mouse squamous cell carcinomas, sequenced on PromethION, were obtained from https://github.com/ugdastider/long_read_paper, and used to visualize *Panx1* transcript isoforms. Figures were generated using the UCSC genome browser (https://genome.ucsc.edu/index.html) (Kent *et al*., 2002).

### Cell surface biotinylation and co-immunoprecipitation

For cell surface biotinylation experiments performed in HEK293T cells expressing different mPANX1 proteins, cells were labeled with 1 mg/ml EZ-Link Sulfo-NHS-LC-Biotin™ (Thermo Fisher Scientific #21335) in PBS for 1 h at 4 °C, 24 h after transfection. The reaction was quenched by incubating the cells in cold PBS containing 100 mM glycine at 4°C for 20 min, then cells were lysed using PBS containing 1% Triton X-100 and a cocktail of protease inhibitors (Sigma-Aldrich #P8340). Biotinylated proteins were collected by incubating cell lysates with streptavidin-agarose beads (Thermo Fisher Scientific #20349) at 4 °C for 2 hours.

For cell surface biotinylation with untransfected and M211-expressing (5.5 µg plasmid in a 10 cm plate, 24 h after transfection) SCC-13 cells, cells were labeled with 0.4 mg/mL EZ-Link™ Sulfo-NHS-SS-Biotin (Thermo Fisher Scientific #A39258) for 20 min at 4°C with gentle shaking. Biotin was quenched with 100 mM glycine-DPBS without Ca^2+^ and Mg^2+^ for 30 min at 4°C with gentle shaking after two quick 100 mM glycine-DPBS without Ca^2+^ and Mg^2+^ rinses. Cells were then lysed with SDS lysis buffer (1% Triton X-100, 0.1% SDS, 1 mM NaF, 1 mM Na_3_VO_4_ in DPBS without Ca^2+^ and Mg^2+^) and protein was quantified via BCA assay. Biotinylated proteins were separated out of lysates (1 mg per sample) by 4°C overnight incubation with 30 µL of pre-washed Pierce™ Streptavidin magnetic beads (Thermo Fisher Scientific #88816). The next day, the beads were washed twice with 2x IP buffer and once with 1x DPBS without Ca^2+^ and Mg^2+^ using the DynaMag™-2 magnet (Invitrogen #12321D), resuspended in 50 µL of 2x Laemmeli buffer and boiled for 5 min before running in parallel with 70 µg of input lysates on a 10% acrylamide gel. E-cadherin and GAPDH were used as positive and negative controls, respectively, for cell surface trafficking.

For co-immunoprecipitation experiments, cells were lysed using 1x IP buffer (1% Triton X-100, 150 mM NaCl, 10 mM Tris, 1 mM EDTA, 1 mM EGTA, 0.5% Nonidet P-40) supplemented with 1 mM NaF, 1 mM Na_3_VO_4_ and half of a Pierce Protease Inhibitor EDTA-free Mini Tablet. Co-immunoprecipitation was performed using 1 mg of Hs578T *PANX1* KO cells co-transfected with PANX1 WT and M211 plasmids (2.75 µg of each plasmid in a 10 cm plate, 24 h after transfection), Pierce™ Protein G magnetic beads (Thermo Fisher Scientific #88847) and the DynaMag™-2 magnet. The manufacturer’s protocol was followed except that 25 µL of beads per sample was used, the lysate and antibody-bound beads were incubated for 30 min at room temperature and antibodies were dissociated from beads by incubating the beads with 20 µL 50 mM glycine elution buffer and 10 µL 2x Laemmli sample buffer (Bio-Rad Laboratories #1610737) supplemented with 10% β-mercaptoethanol for 10 min at 70°C in a T100TM Thermal Cycler (Bio-Rad Laboratories). The following antibodies were used for immunoprecipitation: anti-PANX1 CT-412 10 µg (Penuela *et al*., 2007; Penuela *et al*., 2009), anti-PANX1 NT 1:100 (Cell Signaling Technology #91137, RRID:AB_2800167) and mouse anti-HA 5 µg (Invitrogen #26183, RRID:AB_2533049). Equal amounts of either rabbit or mouse IgG antibodies (Thermo Fisher Scientific #02-6102, #02-6502) were used as pull-down controls. To try to minimize background due to IgG antibodies, blots were probed using the Quick Western Kit (LI-COR Biosciences #926-69100) and the corresponding antibody dilutions outlined above.

### Deglycosylation

To examine whether ATI isoforms of mPANX1 were glycosylated, 50 µg or 60 µg of protein lysates were incubated with glycerol (50%) or 7000 U/mg PNGase F (New England Biolabs #P0704) at 37 °C for 3 h before SDS-PAGE separation. For analysis of M211 glycosylation, 20 µg of lysates from Hs578T *PANX1* KO cells co-transfected with hPANX1 WT and M211 were subjected to 1000 U of PNGase F (New England Biolabs, Inc. #P0704S) at 37°C for 2 h or 1000 U Endo H (New England Biolabs, Inc. #P0702) for 1 h at 37°C. To visualize any shifts in PANX1 banding, lysates from all deglycosylation assays were separated on either 10% or 12% SDS-PAGE gels.

### Electrophysiology

Whole-cell recordings were performed at room temperature using an Axopatch 200B amplifier and pClamp10 software (Molecular Devices). Ramp voltage commands were applied at 7 s intervals, with voltage ranging from −130 mV to 80 mV (0.2 mV/ms) by using the pClamp10 software and a Digidata 1322A digitizer (Molecular Devices). Micropipettes of 3-5 MΩ were pulled from borosilicate glass capillaries (Harvard Apparatus) using a P-97 Micropipette Puller (Sutter Instrument). Bath solution contained 140 mM NaCl, 3 mM KCl, 2 mM MgCl_2_, 2 mM CaCl_2_, 10 mM HEPES and 10 mM glucose (pH 7.3). Internal (pipette) solution was composed of 30 mM tetraethylammonium chloride, 100 mM CsMeSO_4_, 4 mM NaCl, 1 mM MgCl_2_, 0.5 mM CaCl_2_, 10 mM HEPES, 10 mM EGTA, 3 mM ATP-Mg and 0.3 mM GTP-Tris (pH 7.3). CBX-sensitive current was defined as the difference in current at +80 mV, before and after applying 50 μM of CBX (Sigma-Aldrich #C4790) in bath and was normalized to cell capacitance (i.e., current density). Phenylephrine-induced current was determined by the difference in CBX-sensitive current at +80 mV, before and after applying 20 μM of phenylephrine (Sigma-Aldrich #P6126) in bath.

### Statistics

Statistical analyses were performed using GraphPad Prism 10 software. Statistical tests as well as biological replicates are reported in individual figure legends.

## ACKNOWLEDGEMENTS

We thank Dr. Samy Lamouille (Virginia Tech) and Dr. Michael Koval (Emory University) for their feedback and suggestions for the study. This work was supported by Canadian Institutes of Health Research CIHR Project grants (PJT 206190, FRN 153112 to S.P. and 185953 to S.P., M.H. and J.R.) and a United States National Heart, Lung, and Blood Institute grant (P01 HL120840) to D.A.B.

## AUTHOR CONTRIBUTIONS

Conceptualization: B.O’D., M.J.Z., D.J., L.G., S.S., J.W.S., Y-H.C., D.A.B., S.P.; Data Curation: D.S., M.J.Z., Y-H.C.; Formal Analysis: B.O’D., D.S., M.J.Z., S.E.L., L.A.O., J.T., L.G., Y-H.C.; Funding Acquisition: J.J.K., M.H., J.R., D.A.B., S.P.; Investigation: B.O’D., D.S., M.J.Z., D.J., S.E.L., L.A.O., J.T., T.J.F., J.J.K., K.B., S.S., Y-H.C.; Methodology: B.O’D., D.S., M.J.Z., S.E.L., D.J., L.G., J.J.K., J.W.S., Y-H.C., D.A.B., S.P.; Project Administration: B.O’D., D.J., L.G., S.P.; Resources: D.W.L., K.R., J.W.S., M.H., J.R., D.A.B., S.P.; Supervision: B.E.I., D.W.L., K.R., J.W.S., M.H., J.R., D.A.B., S.P.; Visualization: B.O’D., D.S., M.J.Z., J.T., S.E.L., C.VK., L.A.O., Y-H.C.; Writing – original draft: B.O’D., M.J.Z., Y-H.C., S.P.; Writing – review and editing: all authors.

## Abbreviations

ATI: Alternative translation initiation,
CBX: Carbenoxolone,
Cx43/*GJA1*: Connexin 43
GJA1-20k: 20 kDa isoform of connexin 43
PANX: Pannexin
HEK: Human embryonic kidney
hPANX1-FL: Human PANX1 full-length
mPANX1-FL: Mouse PANX1 full-length
mPANX1-25K: 25 kDa isoform of mPANX1
TEV: Tobacco Etch Virus

